# Responses to ligand binding in the bacterial DNA sliding clamp β-clamp manifest in dynamic allosteric effects

**DOI:** 10.1101/2024.06.26.600769

**Authors:** Signe Simonsen, Andreas Prestel, Eva C. Østerlund, Marit Otterlei, Thomas J.D. Jørgensen, Birthe B. Kragelund

## Abstract

The homo-dimeric, ring-shaped bacterial DNA sliding clamp, β-clamp, is a central hub in DNA replication and repair. It interacts with a plethora of proteins via short linear motifs, which bind to the same hydrophobic binding pocket on β-clamp. Although the structure, functions and interactions of β-clamp have been amply studied, less focus has been on understanding its dynamics and how this is influenced by ligand binding. In this work, we have made a backbone nuclear magnetic resonance (NMR) assignment of the 83 kDa dimeric β-clamp and used NMR in combination with hydrogen-deuterium exchange mass spectrometry to scrutinize the dynamics of β-clamp and how different ligands affect this. We found that binding of a small peptide from the polymerase III α subunit affects the dynamics and stability of β-clamp. This effect not only appears locally around the binding pocket, but also globally through dynamic allosteric connections to distant regions of the protein, including the dimer interface. This dissipated dynamic effect is likely a consequence of the binding pocket architecture and may reflect a common mechanism of structural plasticity, where different ligands impose differential responses in the structure and dynamics of β-clamp.

**Highlights:**

- NMR spectroscopy of β-clamp revealed its inherent dynamics
- A peptide ligand from polymerase III stabilizes β-clamp from global unfolding
- Peptide binding causes allosteric effects that manifest in changes in dynamics
- Dynamic changes occur distant to the binding pocket in the dimer interface
- Allostery may be a mechanism for differential responses to interaction partners

**Graphical abstract:** 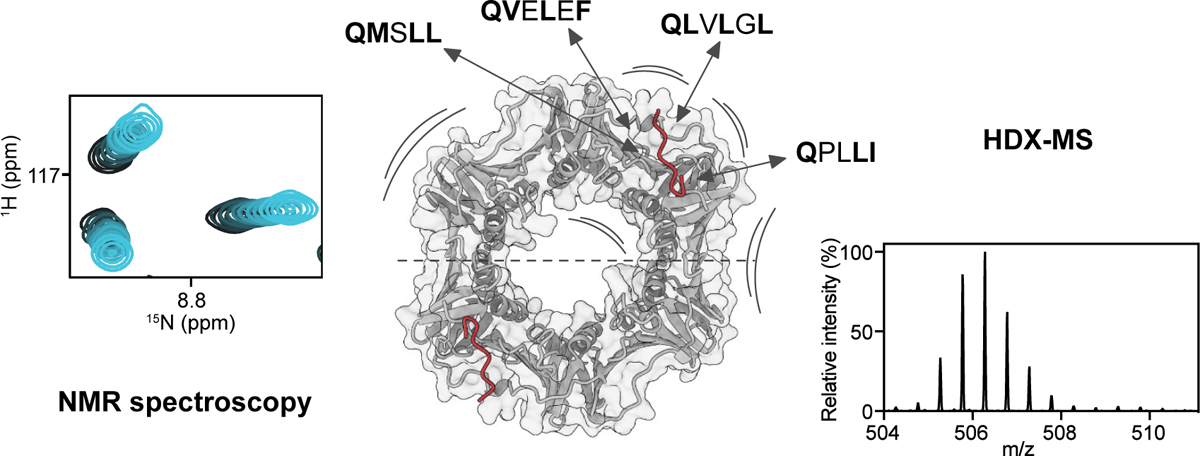

## Introduction

In the bacterium *E. coli*, DNA replication is carried out by a multiprotein complex consisting of a polymerase III complex (pol III core complex), a helicase/primase complex and a clamp loader complex in addition to a DNA sliding clamp, called β-clamp [1,2]. β-clamp is a homo-dimeric, toroidal protein that encircles DNA [3] tethering polymerases to the primer/template junction during DNA synthesis, ensuring processive DNA replication [1,2]. β-clamp is loaded onto DNA in an ATP-dependent crab-claw like motion by the clamp loader complex [4,5] constituting a central hub that interacts with numerous proteins important in translesion synthesis (TLS) [6–8], DNA mismatch repair [9,10] or otherwise involved in DNA homeostasis [10–12].

The structure of each β-clamp subunit consists of three topologically similar domains (I, II and III) connected by interdomain connecting loops (IDCLs) forming a head-to-tail ring-shaped homodimer (**Figure 1A**). The two subunits create an inner channel lined by twelve α-helices and an outer surface composed of 24 β-strands forming antiparallel sheets [13,14]. The many binding partners interact with β-clamp through a short linear motif (SLiM) [15,16] called the clamp binding motif (CBM), which spans five to six residues and is defined as Qφx[L/M][F/L] and Qφx[L/M]x[F/L] (φ = aliphatic, x = any residue) [12] (**Figure 1B**). All known natural β-clamp ligands bind to the same two identical highly conserved binding pockets located between domains II and III (**Figure 1A**). Here, the Gln and the following aliphatic amino acid of the CBM bind to one subsite of the binding pocket, and the two C-terminal hydrophobic residues bind to an adjacent subsite [12]. The binding pocket is overall hydrophobic surrounded by positively charged residues (**Figure 1B**). Although the isolated CBMs bind to β-clamp with micromolar affinity, residues outside the CBM also contribute to affinity [17,18], binding interface [7,11,19] and subsequently to increased lifetime of protein:DNA complexes [20].

**Figure 1:**
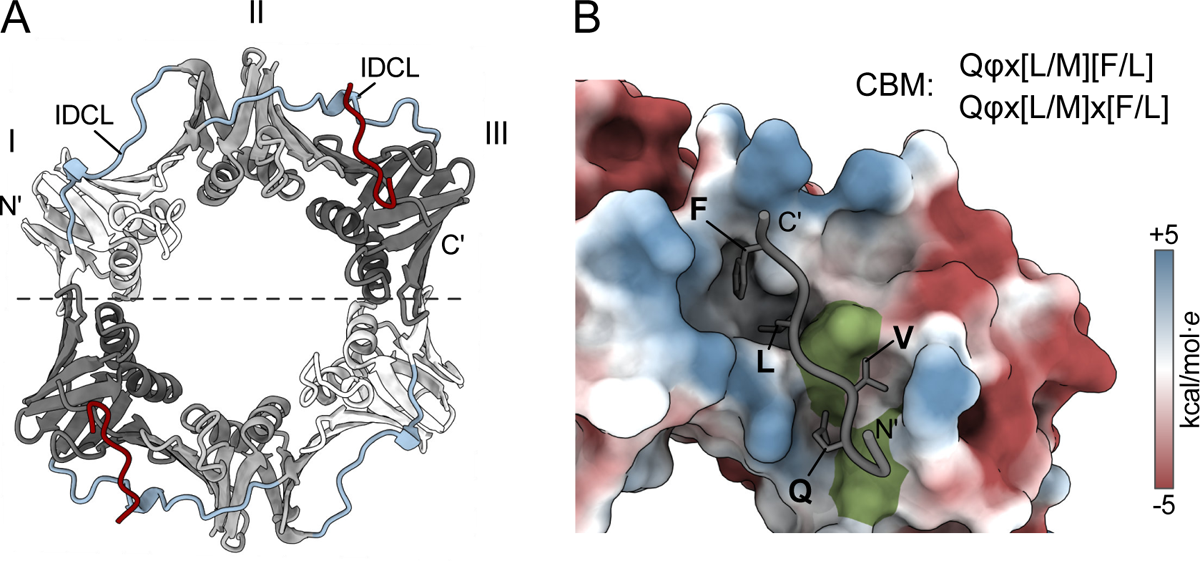
β-clamp structure and binding pocket. A: Structure of *E. coli* β-clamp in complex with a CBM containing peptide from pol III α (PDB ID: 3D1F [30]). β-clamp domains I, II and III are shown in white, light grey and dark grey, respectively, and IDCLs in blue and the pol III peptide in red. B: The β-clamp binding pocket with the electrostatic potential computed in ChimeraX. The pol III peptide is shown in grey with key SLiM-interacting residues highlighted as sticks. Two methionine residues of β-clamp, Met362 and Met364 laying within the binding pocket are coloured green.

β-clamp is a stable dimer in solution with an estimated *K*_D_ in the low picomolar range in the absence of DNA [21,22] and a half-life for spontaneous DNA dissociation of ∼1-2 hours [21,23,24]. Still, studies have found β-clamp to be dynamic in solution. While hydrogen-deuterium exchange mass spectrometry (HDX-MS) studies concluded that domain I is highly dynamic [25,26], a high-resolution crystal structure of β-clamp showed increased B-factors in domains II and III [14]. It was also suggested that the β-clamp dimer can open spontaneously existing in an open-closed equilibrium [19,25], while several other studies have found that the major state of β-clamp is a closed dimer [19,27–29]. Thus, there is an apparent inconsistency regarding how and in what ways β-clamp is dynamic, possibly reflecting different conditions or methodology. Still, very little is known about the dynamic properties of β-clamp and how this is influenced by ligand binding.

Here we combine NMR spectroscopy with HDX-MS to address the conformational dynamics of β-clamp and we scrutinize how binding of CBM-carrying ligands affects this. With the dynamics mapped on several timescales, we found that peptide binding influences both fast and slow-timescale β-clamp dynamics, conformational stability, and a tendency to self-associate at high concentrations. Not surprisingly, regions in domains II and III that entail the protein binding pockets become stabilized upon ligand binding. Remarkably, ligand binding allosterically affects the dimer interface far from the binding pockets, likely facilitated by a unique architecture of the binding pocket. This structural and dynamic adaptability of β-clamp could have implications for its function as a hub and allows β-clamp to interact with a wide variety of protein binding partners though the same shared site. With variations in the CBMs, different ligands may affect the dynamics of β-clamp differentially and lead to different functional outcomes.

## Results

### Strategy to circumvent unfavourable dynamics for backbone assignment of β-clamp

To investigate the dynamics and interactions of the bacterial DNA sliding clamp, we made a backbone assignment of recombinantly produced β-clamp. Due to its very large size (83 kDa) β-clamp was expressed in ∼97% deuterated minimal media supplemented with ^2^H, ^13^C_6_ labelled glucose and ^15^N labelled NH_4_Cl. Expression and purification of ^2^H, ^13^C, ^15^N β-clamp yielded ∼20-30 mg pure protein per 0.5 L expression media.

Although the TROSY-HSQC of ^2^H, ^13^C, ^15^N labelled β-clamp showed nice dispersion and fair intensities of the peaks (**Figure S1A**), the resonance assignment proved challenging. HNCA and HNCO experiments yielded decent spectra, however the spectral quality of HNCACB spectra were inadequate for assignment. Instead, modified HNCA experiments called gradient optimized CO decoupling pulse (GOODCOP) and beta/alpha decoupling pulse (BADCOP) were implemented [31]. These spectra provide indirect information on the C^β^ chemical shift via selective decoupling and help resolve ambiguities of sequentially matching peaks in the HNCA spectra during assignment. Furthermore, 3D-^15^N-TROSY-NOESY [32] spectra were obtained that together with crystal structures of β-clamp aided assignment of spatially close residues in the β-clamp structure. Despite these useful NMR experiments, assignments of the β-clamp stalled after approximately 50-60% of the protein backbone resonances. The main reason was low spectral quality of the 3D spectra, resulting in many missing peaks in the HNCA spectra.

Because β-clamp tended to precipitate over time at high concentrations at 37°C, we aimed to stabilize the protein by adding a ligand. Remarkably, addition of a small eight-residue peptide harbouring the C-terminal CBM of pol III α (Ac-EQVELEFD, hereafter denoted as pol III) improved the quality of the TROSY-^1^H, ^15^N-HSQC spectrum, but was accompanied with substantial movement of many peaks (**Figure S1B**). Importantly, in the presence of pol III, the quality of the 3D-spectra markedly improved; HNCACB spectra yielded distinct peaks of both C^α^ and C^β^ nuclei and the minor peaks missing in the HCNA/GOODCOP/BADCOP spectra of unbound β-clamp became intense (**Figure S2**). The resonances of pol III-bound β-clamp were therefore assigned using HNCA, HNCACB, GOODCOP, BADCOP and TROSY-NOESY experiments. To track the peaks of the pol III bound β-clamp back to the peaks of unbound β-clamp, a TROSY-^1^H, ^15^N-HSQC titration was made (Figure 2A). To ensure a correct match between the bound and unbound peaks, especially in parts of the spectrum with substantial spectral overlap in the TROSY-HSQC, C^α^ chemical shifts and NOESY connections were compared.

**Figure 2:**
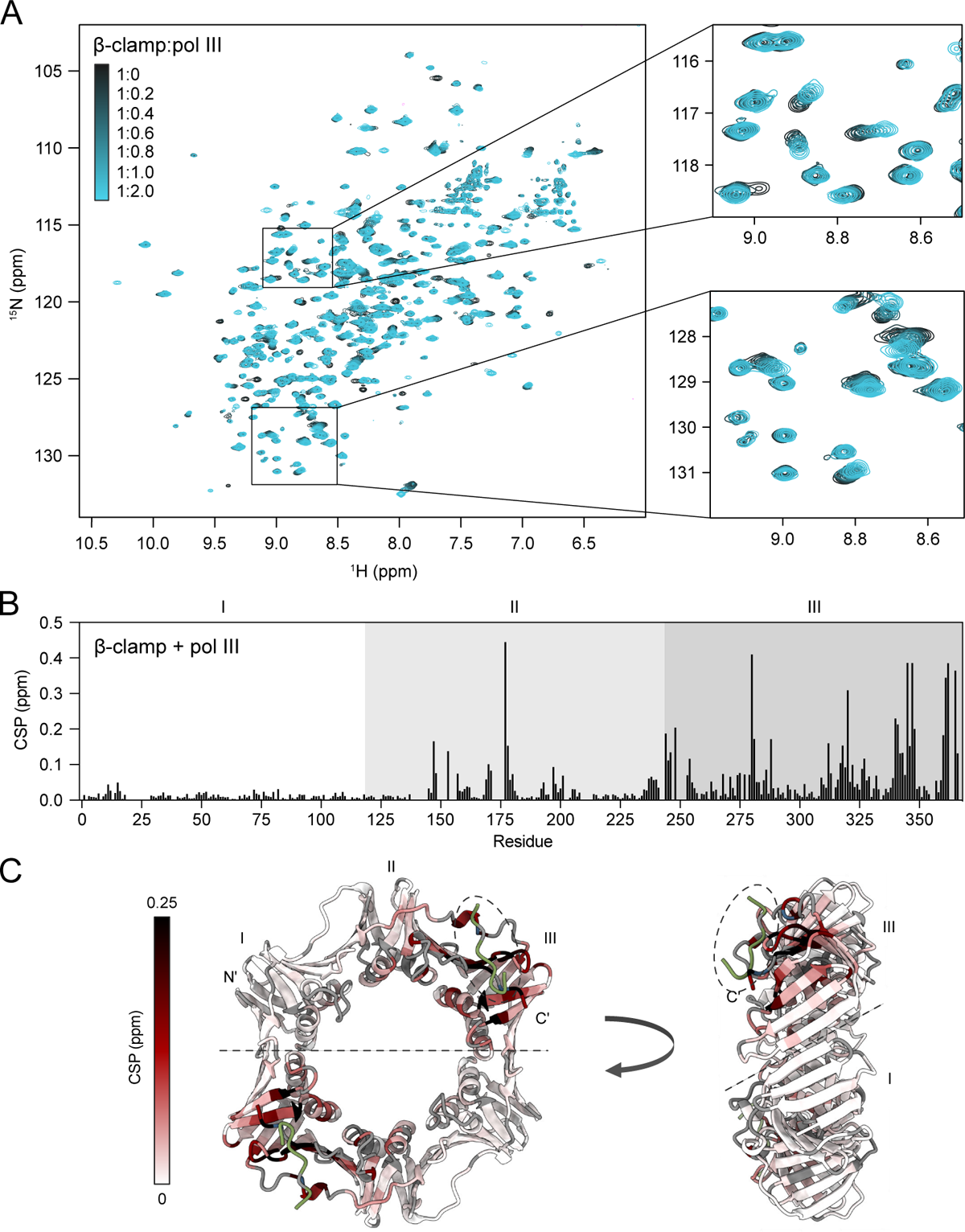
Peptide binding impacts the dynamics of the protein binding pocket on β-clamp. A: ^1^H, ^15^N TROSY-HSQC titration of a ^2^H, ^13^C, ^15^N β-clamp with pol III peptide. B: Chemical shift perturbations between unbound β-clamp and β-clamp:pol III (1:2) for all assigned residues. C: Chemical shift perturbations upon pol III binding mapped onto the structure of β-clamp in complex with a pol III peptide (PDB ID: 3D1F [30]). The peptide is in green and highlighted with a dashed circle. Unassigned residues are coloured grey.

The strategy resulted in the assignment of, 86% of the unbound β-clamp and 89% of the pol III bound β-clamp (excluding prolines and the His_6_-tag). Unassigned residues mainly locate to flexible loops or small turns between connecting β-strands (**Figure S3**). Their peaks were likely missing due to structural heterogeneity on the intermediate exchange regime or due to fast proton exchange with the solvent at the high temperature. A few residues were only assigned in the β-clamp + pol III assignment. Met364, which interacts closely with pol III [30], was only assigned in free β-clamp. Although Ala147, Tyr153 and Gly157 were not assigned in the free β-clamp, they were visible in the titration spectra with pol III.

### Ligand binding induces global changes in β-clamp dynamics

The major spectral changes induced by binding of the eight-residue peptide indicated a global change in the structure and/or dynamics of β-clamp. We therefore mapped the chemical shift perturbations (CSPs) for each residue (Figure 2B). The CSPs mainly located to domain III with domain II less affected, and only minor perturbations in domain I. Mapping the CSPs onto the structure of β-clamp clarified that the largest CSPs were from residues located in or around the binding pocket (Figure 2C). Although we expect residues interacting with the peptide to display large CSPs, residues further away from the binding site were also highly affected suggesting that ligand binding causes a change in structure and/or dynamics of β-clamp. Secondary chemical shifts (SCSs) which report on secondary structures were generally consistent with the secondary structure content of the β-clamp crystal structures (**Figure S4**). Yet, the SCSs of β-clamp with and without the addition of pol III were overall very similar, except for the extreme C-terminal residues directly involved in the interaction with pol III (**Figure S4**). This suggest that changes upon ligand binding do not occur on the secondary structural level. Furthermore, β-clamp has been crystallized in the absence [13,14] and presence of a similar nine-residue peptide of the pol III α C-terminal CBM [30]. These crystal structures align with an RMSD of C^α^ atoms of only 0.95 Å. Thus, differences in the 3D structures cannot explain the large spectral changes we observe upon ligand binding (**Figure S5**), suggesting them to originate from changed dynamics.

To address whether and how ligand binding affects the dynamics of β-clamp, peak intensity ratios from the unbound and pol III-bound β-clamp were mapped for each residue (**Figure S6**). An overall increase in intensity of the peptide-bound β-clamp was in line with the enhanced quality of the NMR spectra (Figure 3A and B). Notably, peak intensities of three residues located in the loop formed by residues 145-157 increased markedly, indicating that ligand binding stabilizes this loop. This is also the loop where most residues are either not assigned or assigned only in the pol III-bound state. Some residues also show decreased peak intensities in the ligand bound state, but these are mostly located at the binding interface (Figure 3A and B) and likely experience faster relaxation due to dipolar coupling to protons of the pol III peptide.

**Figure 3:**
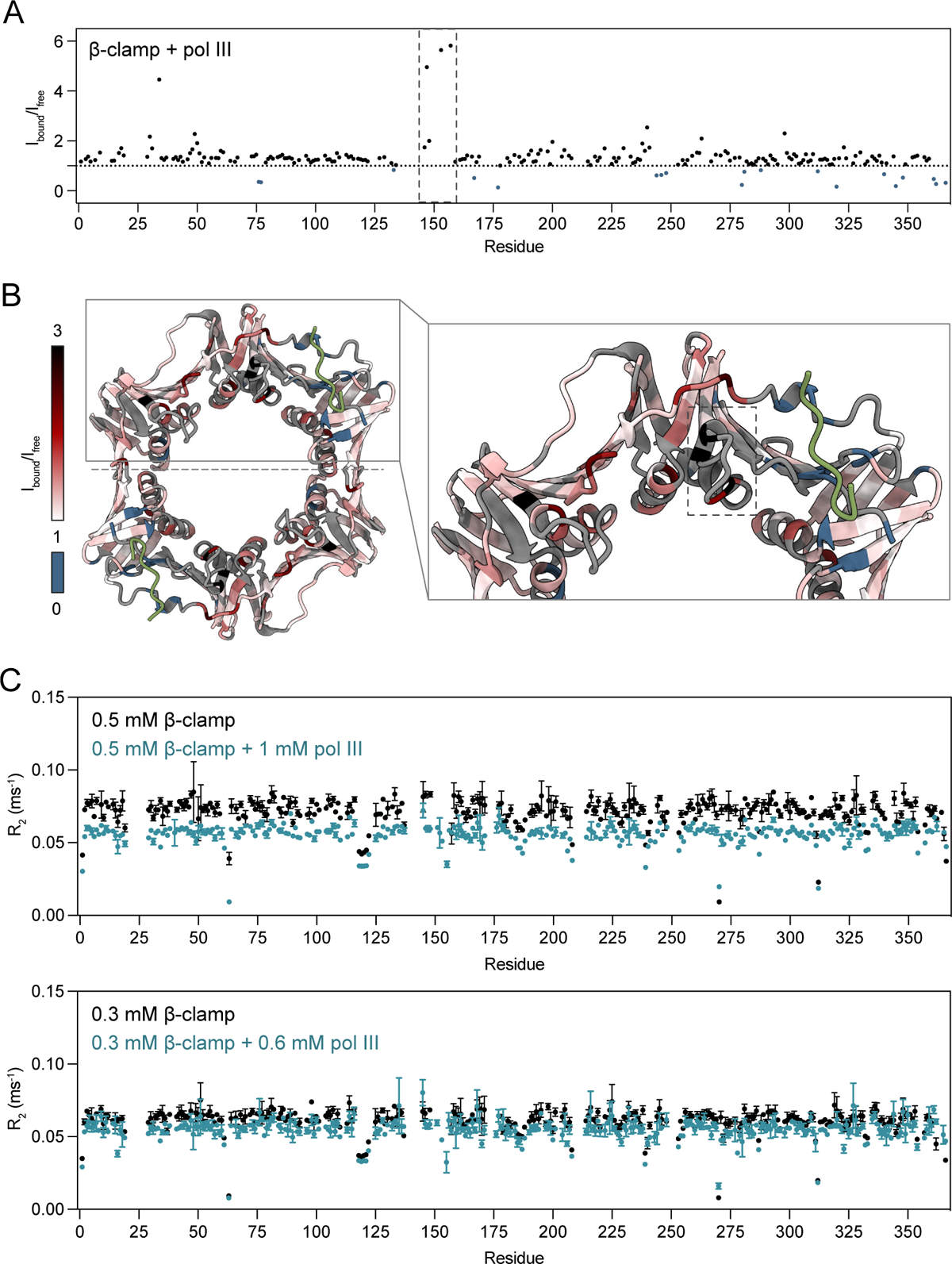
Peptide binding stabilizes β-clamp dynamics and reduces self-association. A: Peak intensity ratios between pol III bound and unbound β-clamp. B: Intensity ratios mapped onto the β-clamp structure in complex with a pol III peptide (green) (PDB ID: 3D1F [30]). Residues for which peak intensities decrease upon peptide biding are coloured blue. Residues with the largest increase in peak intensities are highlighted by a dashed square. C: *R*_2_ values of β-clamp in the absence (black) and presence (blue) of two-fold molar excess of pol III. Top and bottom plots are *R*_2_ values of β-clamp at concentrations of 0.5 and 0.3 mM, respectively.

Because the peak intensities are related to the spin-spin relaxation, we recorded *R*_2_ relaxation rates on β-clamp in the presence and absence of pol III (Figure 3C). *R*_2_ values were generally high and uniform, characteristic of a large, folded protein. However, at a concentration of 0.5 mM β-clamp, we noticed that the unbound β-clamp had overall larger *R*_2_ values compared to the pol III-bound protein (Figure 3C). Since the overall *R*_2_ is related to the overall size of the molecule, we speculated whether β-clamp tends to self-associate at high concentrations and repeated the relaxation measurements at lower concentrations. Here, the pol III-bound β-clamp *R*_2_ values were essentially unaffected by concentration changes, while *R*_2_ values of free β-clamp decreased with decreasing concentration, approaching those of the ligand-bound state. These results suggest that β-clamp tend to self-associate at high concentrations and that ligand binding abolishes this, explaining the improvement in spectral quality.

### Global changes in chemical shifts are a universal effect of peptide binding to β-clamp

Next, we explored if the ligand-induced global effect on the chemical shifts occurred upon binding of other peptide ligands and recorded ^1^H, ^15^N TROSY-HSQCs of β-clamp in complex with peptides from polymerase IV (pol IV) and from two proteins involved in mismatch repair, MutS and MutL (**Table 1**, **Figure S7**). For quantification, we designed the peptides as 15-16-mers including an N-terminal Trp. For better comparison, a similar, long version of the pol III peptide with a Trp was included. These longer peptide variants are denoted with an asterisk. To investigate how the flanking region of the long pol III* affect the interaction with β-clamp, we compared the binding of the 16-mer pol III* to that of the eight-mer pol III. ^1^H, ^15^N TROSY-HSQCs and isothermal titration calorimetry (ITC) showed that the flanking region do contribute to the interaction by forming additional contacts to β-clamp, without largely affecting the binding affinity (**Figure S8**).

**Table 1:**
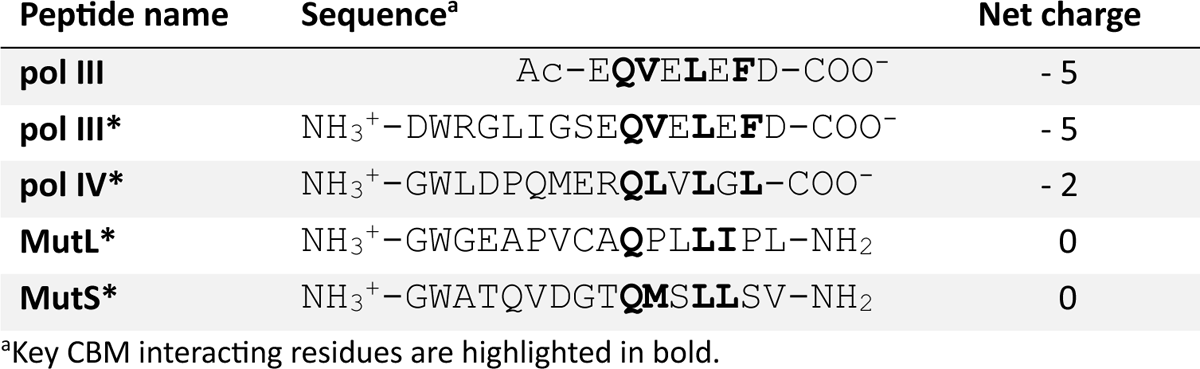
Peptide sequences and properties.

Comparing the NMR spectra of the complexes revealed that the global changes were also induced by binding of the pol IV* peptide (Figure 4). Titrations with the pol III* and pol IV* peptides showed large effects on β-clamp residues of domain II and III which surround the binding pocket. Notably there were slight differences; while the CSPs of domain I were very similar for the two peptides, pol IV* induced slightly larger CSPs in domain II and especially in the interface between domains II and III. Conversely, pol III* generally induced larger CSPs in domain III (Figure 4). This is despite the similar affinities of the pol III and pol IV CBMs [12,30,33,34]. The differences could be caused merely by the different chemical nature of the residues, where e.g., pol III* is more negatively charged compared to pol IV*, or because the flanking regions outside the binding motifs interact differently with β-clamp. However, the widespread effect points towards a differential effect in the interaction of the CBM binding residues with the binding pocket, causing different impact on the structural conformation and dynamics of the β-clamp.

**Figure 4:**
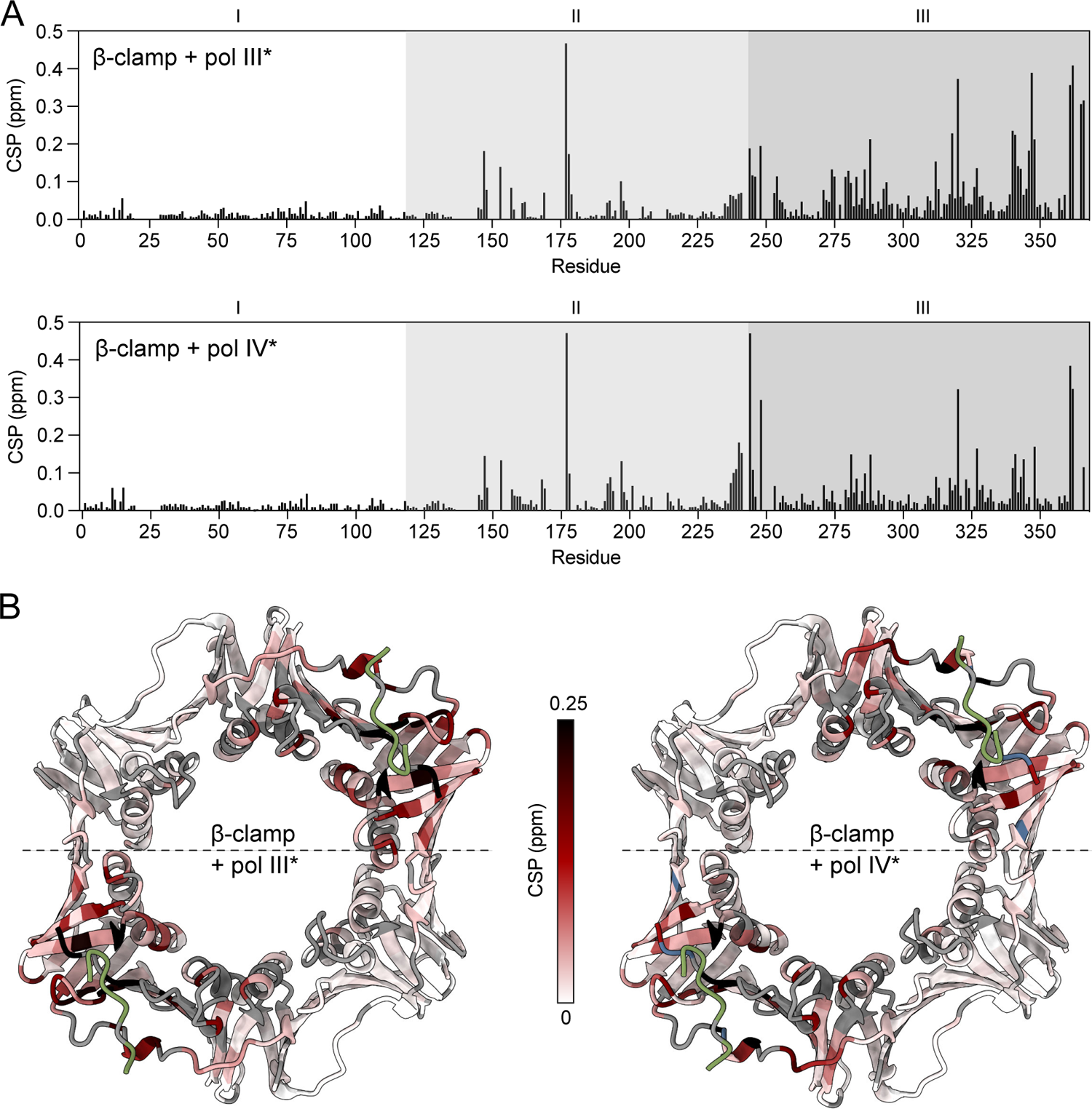
pol IV* binding causes different global changes on the chemical shifts compared to pol III*. A: CSPs derived from ^1^H, ^15^N TROSY-HSQCs of ^2^H, ^13^C, ^15^N labelled β-clamp in complex with pol III* (top) and pol IV* (bottom). B: CSPs mapped onto the structure of β-clamp in complex with pol III peptide (PDB ID: 3D1F [30]). Unassigned residues are coloured grey. Peaks that were untraceable due to intermediate exchange are coloured blue.

The CBM of MutS also induced CSPs for many of the same residues as pol III* and pol IV*, however, their amplitudes were smaller (**Figure S7**). To our knowledge, there are no reported affinities on MutS [9,33] and it may be that the β-clamp:MutS* interaction is not completely saturated, explaining the smaller effect. There are also no reported affinities on MutL, but the affinity is likely very low [9,35]. Although we would expect the peaks to move in fast exchange at such low affinity, the binding of the CBM of MutL appeared to cause substantial line broadening (**Figure S7**). Thus, MutL* may bind β-clamp differently than the other CBM peptides. The CBM of MutL differs from the canonical CBM in that it contains a Pro at the second position and a Leu/Ile combination at position 4 and 5 [12]. Crystal structures of the complex between MutL and β-clamp also show that the CBM bind differently in the β-clamp binding pocket, likely caused by the proline [9,12]. It is possible that the line broadening therefore is caused by structural heterogeneity within the binding pocket [34].

### β-clamp experiences fast and slow hydrogen-exchange

To further investigate the dynamic properties of β-clamp and the impact from ligand binding, we performed HDX-MS on free β-clamp and β-clamp in complex with pol III. At first, the global deuterium uptake of intact β-clamp was measured. A non-deuterated control of β-clamp yielded an experimental mass of 41466 Da (Figure 5A), corresponding well with the theoretical mass of 41466.07 Da of one subunit of the His_6_-tagged β-clamp with the N-terminal Met removed. We noticed two additional peaks each with a mass increase of 16 Da, suggesting β-clamp to be partially oxidized carrying up to two oxidations per subunit (Figure 5A).

**Figure 5:**
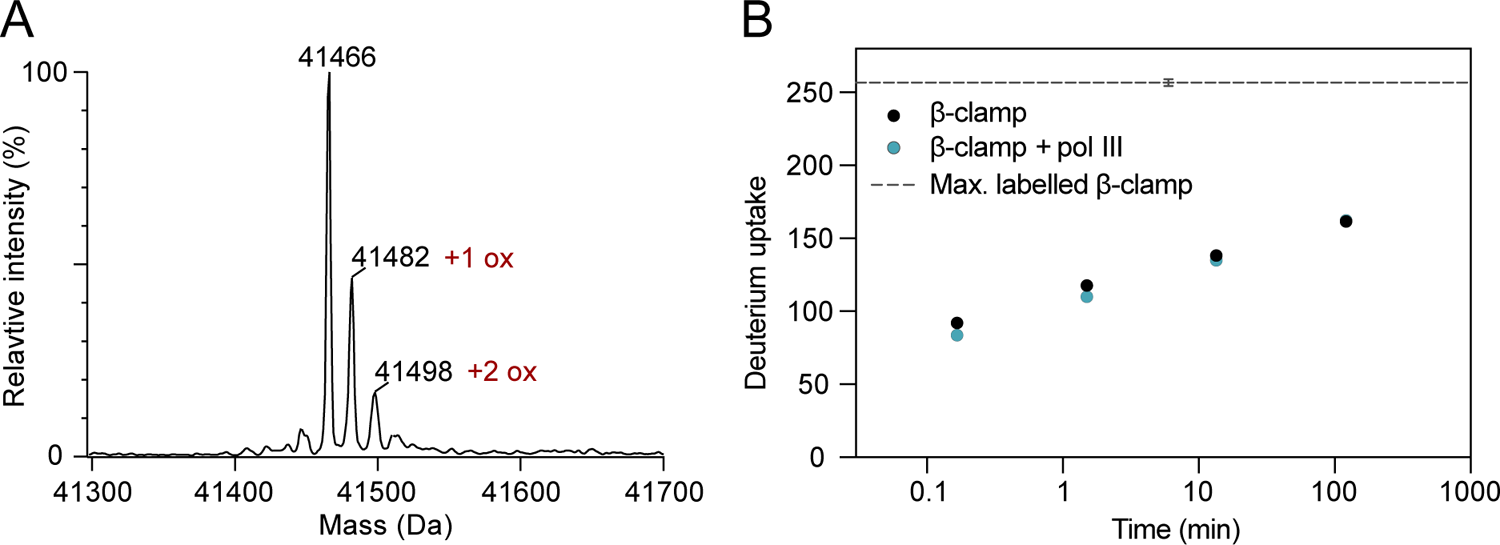
Global HDX-MS analysis reveal a weak protection against exchange in β-clamp upon pol III binding. A: Deconvoluted ESI-MS spectrum obtained from β-clamp in H_2_O-buffer. B: Deuterium uptake of free β-clamp and β-clamp in complex with pol III peptide after incubation for 10 s, 1.5 min, 13.5 min, and 121.5 min in deuterated PBS buffer at 37°C, pD 7.4. The deuterium uptake of the maximally labelled β-clamp sample is indicated by a grey dashed line. The data points are an average of triplicate values, error bars show ±SD, but most of them are too small to be visible in this plot.

During HDX-MS analysis, an inevitable loss of deuterium will occur due to back-exchange with the protonated solvents. To measure the experimental maximum deuterium uptake of β-clamp a maximally labelled control was made. Here, β-clamp was diluted 10-fold in 6M guanidinium deuterochloride (GdnDCl) and incubated for 2 hours resulting in a deuterium uptake of 257 ± 3 Da. Since the theoretical maximum uptake of β-clamp in 90% D_2_O is 317 Da, the back-exchange of β-clamp in the global HDX-MS analyses is 19%.

To probe the effect of pol III on the global deuterium uptake, β-clamp was diluted 10-fold into deuterated buffer to initiate H-to-D exchange in the presence and absence of pol III. After various periods of deuteration, the exchange reaction was quenched by acidification and cooling followed by mass measurement. The result of the global uptake analysis showed that within the first 10 s, there was a fast exchange of the amide protons with an average deuterium uptake of 92 ± 0 Da in unbound β-clamp and 84 ± 2 Da in pol III-bound β-clamp (Figure 5B). After approximately two hours, the deuterium uptake reached 162 ± 2 Da for both samples, which was still far from the maximally labelled β-clamp, i.e., 257 Da. Even after more than four hours, the deuterium uptake only reached 176 Da for free β-clamp and 173 Da for pol III-bound β-clamp (**Figure S9**). The hydrogen-deuterium exchange is therefore defined by a fast exchange process, likely occurring in the unprotected turns and flexible loops in β-clamp, followed by a slower exchange process where a substantial part of β-clamp remains unexchanged after several hours. A weak, yet significant protection induced by binding of pol III is evident by the lower D-uptake of pol III-bound β-clamp vs. free β-clamp at the early time points. As the protection is weak, this difference in deuterium uptake disappears after prolonged deuteration. This suggested that the overall exchange process is similar for the bound and unbound β-clamp.

### Methionine oxidation in the binding pocket affects pol III binding

By local HDX-MS analysis of proteolyzed β-clamp, we identified the methionine oxidations to be located at the extreme C-terminus. Peptide 359-366 was identified as non-oxidized and as oxidized (+16 Da) at either Met362 or Met364. Both methionine residues are located within the protein binding pocket (Figure 1B). Deuterium uptake plots of peptide 359-366 clearly showed that a protection induced by pol III binding at 10 seconds in the non-oxidized β-clamp was almost completely abolished when Met362 was oxidized (Figure 6A). Oxidation of Met364 had the same effect (**Figure S10**). This suggests that oxidation of methionine residues in the binding pocket decreases pol III affinity. We also noticed that upon NMR titrations of β-clamp with various peptide ligands, two populations of the C-terminal Leu366 were present (Figure 6B). Thus, a small population of what appears to be the unbound β-clamp state persists until it disappears with excess amount of ligand. This small population may correspond to the oxidized protein, and further supports different ligand affinity between non-oxidized and oxidized forms of β-clamp. Depending on the degree of oxidation, these observations could have large implications when studying β-clamp.

**Figure 6:**
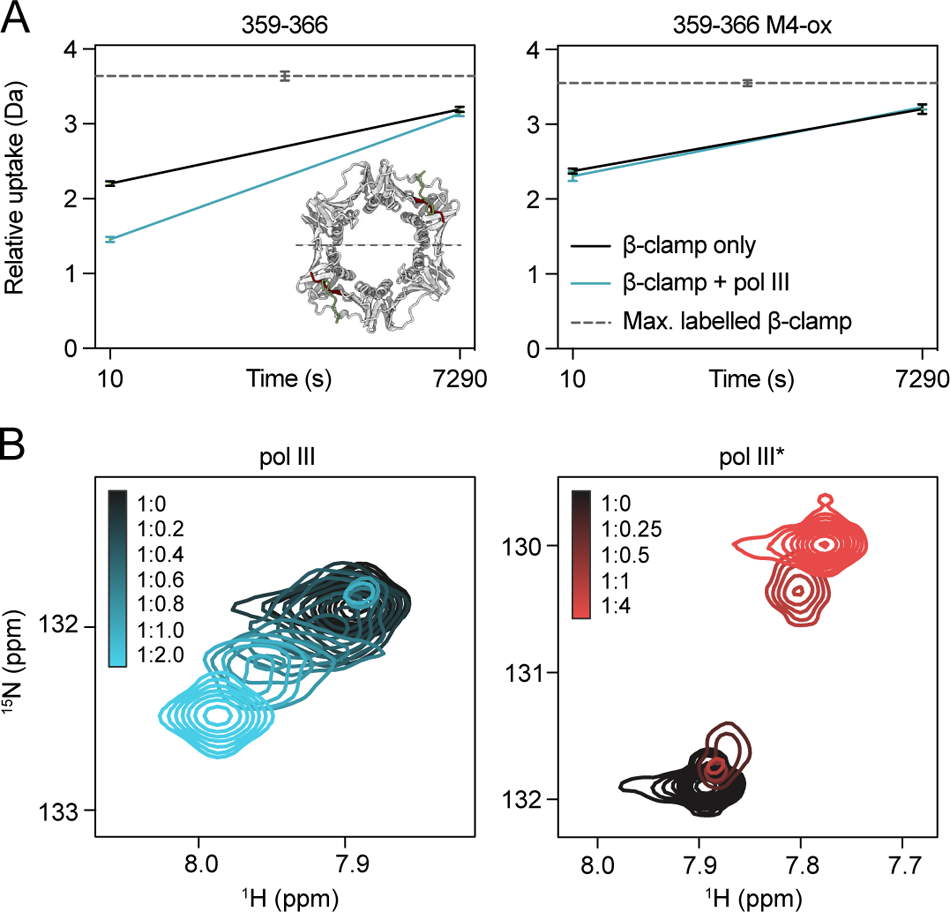
Methionine oxidation in the β-clamp binding pocket impacts binding. A: Local HDX-MS deuterium uptake plots of peptide 359-366 in its non-oxidized form (left) and with oxidation of Met362 (right). B: Zoom on the C-terminal Leu366 from the ^1^H, ^15^N TROSY-HSQC titrations of ^2^H, ^13^C, ^15^N β-clamp with addition of pol III (left) and pol III* (right).

### Peptide binding stabilises β-clamp against global unfolding

We discovered serendipitously by HDX-MS that β-clamp slowly undergoes global unfolding in 4M urea, and that the unfolding kinetics is readily monitored by the emergence of bimodal peaks in the global uptake plots of intact β-clamp, corresponding to two different populations. The low mass peak represents a population of folded β-clamp with a deuterium uptake comparable to that observed without urea, while the high mass peak represents the maximally labelled globally unfolded β-clamp. We furthermore noticed that the two states were differently populated in the presence and absence of pol III, where pol III suppressed the population of the unfolded state relative to the folded state. This led to the hypothesis that pol III may impact the stability of β-clamp. We therefore designed a urea experiment, where β-clamp with and without pol III were diluted 10-fold in 4M deuterated urea and the reaction quenched at different time points. Again, the MS spectra showed two peaks corresponding to a fully deuterated protein and a partially deuterated protein (Figure 7A). Data processing and quantification of the two states revealed that the unfolding reaction is time dependent and slow (**Figure S11**). After incubation for only 10 seconds, the fully unfolded population was not observed. Over time, the population of the unfolded, fully deuterated state gradually increased until the entire population was fully unfolded after 24 hours (Figure 7B). The unfolded population increased by 14.1% and 11.6% per hour for unbound β-clamp and pol III-bound β-clamp, respectively. At the time points between 1-4 hours, where both states are present, there was a significant population difference between free β-clamp and pol III-bound β-clamp, where the fully unfolded state was less populated in the pol III-bound protein samples. Thus, pol III has an overall stabilizing effect on the slow unfolding of the β-clamp in the presence of urea.

**Figure 7:**
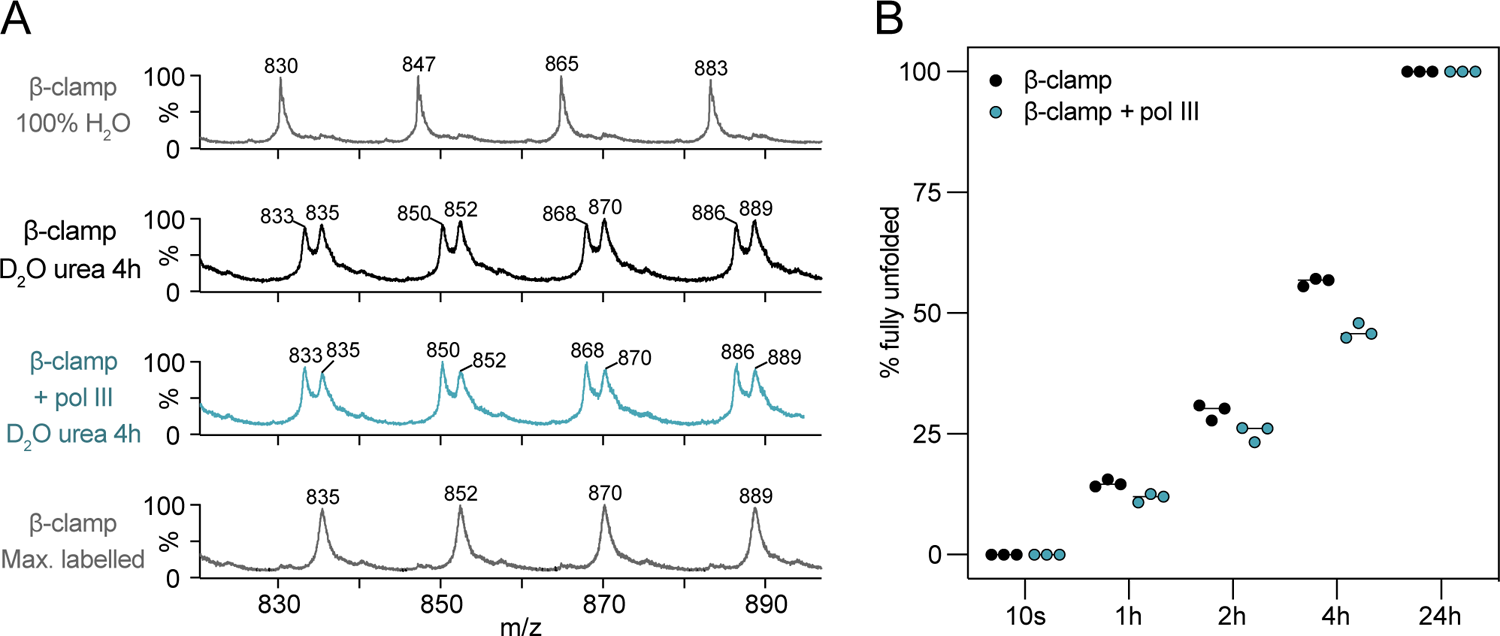
Pol III peptide stabilizes β-clamp against global unfolding. A: ESI-MS spectra showing the two populations present in the samples of free β-clamp and pol III bound β-clamp incubated in 4M deuterated urea at 25°C for 4 hours (two middle panels). Top panel is the non-deuterated control (in H_2_O PBS buffer, pH 7.4), and the bottom panel is the maximally labelled control (6M GdnDCl, 37°C, 2 hours). B: Fraction (%) of the fully deuterated and unfolded state for free β-clamp and pol III bound β-clamp at different time points. Unpaired, two-tailed t-test for significance between unbound and pol III bound samples; 1 hour: p-value = 0.0116 (*); 2 hours: p-value = 0.0317 (*); 4 hours: p-value = 0.0006 (***).

To test if the unfolding by urea was reversible or irreversible, we made a pulse-labelling experiment. Here, β-clamp with and without pol III were incubated in non-deuterated urea. After two hours, the samples were diluted into deuterated urea and incubated for one minute before the exchange reaction was quenched, allowing readily exchangeable hydrogens to exchange with deuterium. If the unfolding by urea was reversible, we would expect the unfolded state to be negligible. Yet, we saw a substantial amount of the fully deuterated state, indicating an irreversible unfolding process (**Figure S11**).

### Pol III binding stabilizes the binding pocket and impose allosteric effects at distant sites

To investigate the β-clamp dynamics in the absence of ligand, we analysed the local deuterium uptake. First, peptic peptides of β-clamp were identified, and their deuterium uptake subsequently measured (the peptides and their deuterium uptake are listed in Supplementary Table S1, sequence coverage map shown in **Figure S12** and example spectra shown in **Figure S13)**. The N-terminal His_6_-tag experienced an abnormal amount of back-exchange (**Table S1**), as previously noted [36,37], and was excluded from the analysis. The average amount of back-exchange for all remaining peptides was 36%.

Representative peptides from β-clamp and their individual HDX in the absence of pol III were mapped onto the β-clamp structure (Figure 8A). To increase the resolution, the peptides were selected to cover as much of the β-clamp structure with as short peptides as possible. The average deuterium uptake for these selected peptides was 45% after 10 seconds and 76% after 2 hours. Peptides with more than 50% deuterium uptake after 10 seconds were all located in the IDCLs, in the very C-terminus or in other flexible loops or linkers, whereas peptides from structured parts of β-clamp remained relatively protected. The same was true after 2 hours, where many of the forementioned loops had completely exchanged with deuterium, while structured parts generally were more protected. The overall HDX in the three domains of β-clamp did not vary greatly. Notably, after 2 hours, peptides 7-16 and 317-325 displayed high deuterium incorporation compared to the average (97% and 89%, respectively) despite not covering any loops (**Figure S14**), hinting that these parts of the structure are highly dynamic compared to other structured regions of β-clamp. Interestingly, the dimer interface appeared to be one of the most stable parts of the structure, with only 13% (10 s) and 55% (2h) uptake for peptide 98-111, and 23% (10 s) and 64% (2h) for peptide 284-306 (**Figure S14**).

**Figure 8:**
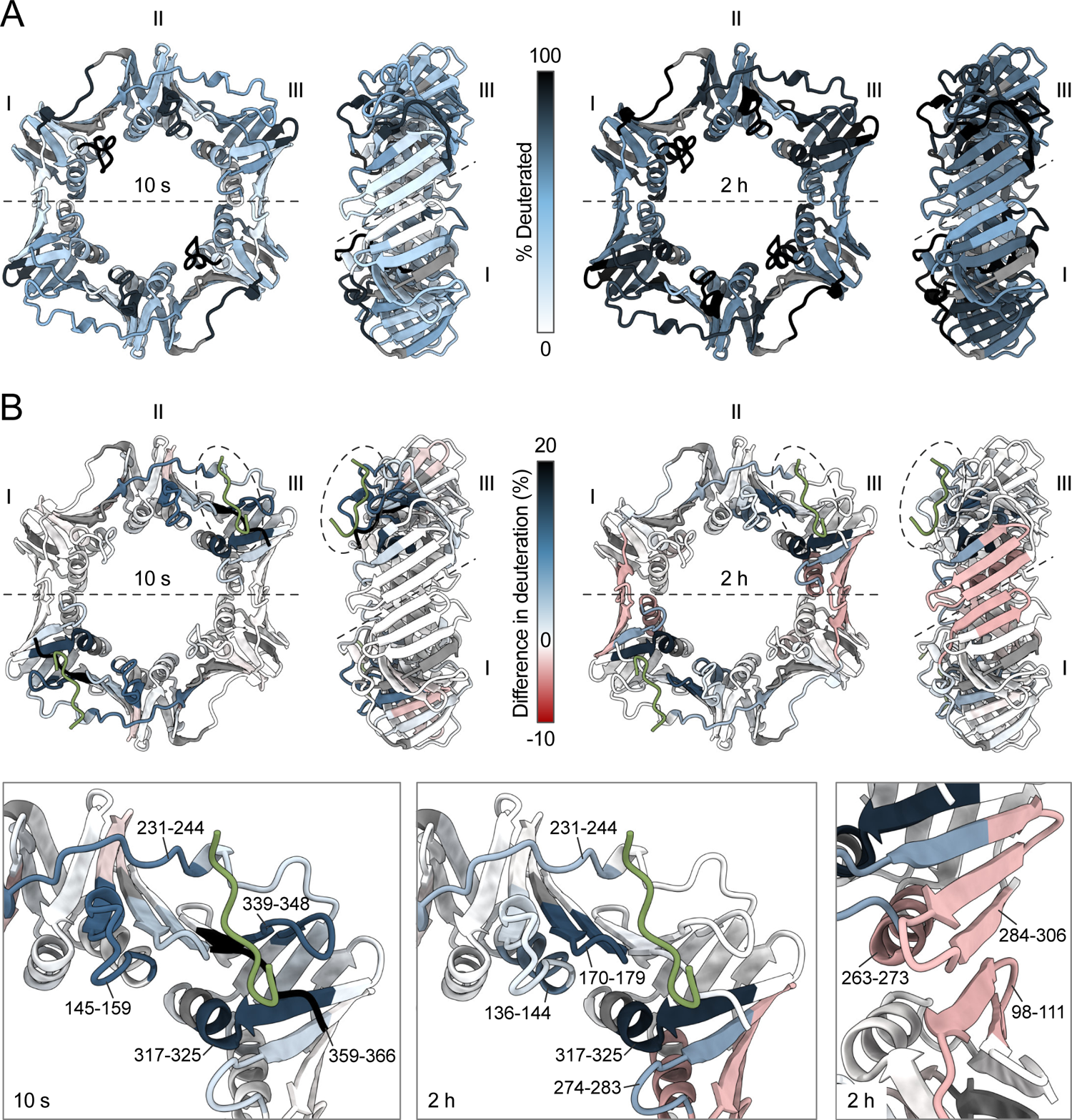
Hydrogen-deuterium exchange mapping shows stabilization of the binding pocket and allosteric effects at distant sites upon ligand binding. PDB ID: 3D1F [30]. pol III is coloured green and unassigned peptides are coloured grey. A: Overall hydrogen-deuterium exchange of selected peptides from β-clamp in the absence of pol III mapped onto the structure of β-clamp. B: The difference in deuteration (%) between free β-clamp and β-clamp + pol III, where positive values (blue) is the result of a lower deuterium uptake in the presence of pol III and negative values (red) is the result of a higher deuterium uptake in the presence of pol III. Bottom panel shows zooms of the protein binding pocket after 10 seconds (left), 2 hours (middle) and of the dimer interface after 2 hours (right).

To further address how pol III affects the dynamics and stability of β-clamp, we compared the deuterium uptake of β-clamp in the absence and presence of pol III and illustrated it on the β-clamp structure (Figure 8B). The difference in %-uptake for free β-clamp and pol III-bound β-clamp for all peptides is also listed in **Table S1** and uptake plots for peptides with a difference in protection in unbound and pol III-bound β-clamp shown in **Figure S15**. After 10 seconds, a substantial protection of peptides that form the binding pocket was observed. Most prominently, peptides 145-159, 231-244, 317-325, 339-348, and especially 359-366, gained protection in the presence of pol III (Figure 8). This is not unexpected, since the pol III peptide interacts with residues within these sequences [30]. While the C-terminal peptide (359-366) is almost completely buried upon binding pol III, as evident from the crystal structure [30], other peptides, such as 145-159 where only side chains interact, likely get protected because the interactions stabilize the dynamics of this loop.

Ligand binding induces weak protection in some peptides that are part of the binding pocket (e.g., peptide 359-366), while other peptides that are also part of the binding pockets are more strongly protected as evidenced by decreased deuterium incorporation in the presence of pol III even after 2 hours, e.g., peptide 170-179 and 317-325 (Figure 8B). Given a β-clamp concentration of 5 μM, a pol III concentration of 50 μM and an approximate *K*_D_ of 3.6 μM (**Figure S8**), the calculated fraction of bound β-clamp at any given time is ∼93%. With an affinity in the micromolar range and a fast exchange between bound and unbound β-clamp seen on the NMR timescale, the peptide would be expected to bind and unbind numerous times within 2 hours. This likely explains why the peptide 359-366, which lies directly in the binding pocket, did not exhibit a reduced deuterium uptake after 2 h in the presence pol III. Intriguingly, strong protection was also observed in peptides that are remote to the binding site, e.g. peptides 136-144, 231-244 and 274-283, indicating that a long-term stabilization manifests in other parts of the β-clamp structure that are not involved in direct interaction with pol III. Thus, combined these data suggest that ligand binding to β-clamp has allosteric effects.

Unexpectedly, the peptides from domain I and III located in the β-strands of the dimer interface (peptides 98-111 and 284-306) as well as the α-helix at the dimer interface of domain III (peptide 263-273) exhibited a slightly higher deuterium uptake in the presence of pol III (Figure 8B). As pol III stabilizes β-clamp against unfolding (Figure 7), we would expect that the dimer interface had a decreased deuterium uptake in the pol III bound state. One possible solution to this conundrum may be that the interaction with pol III represents a noncanonical scenario, in which part of the ligand-binding energy causes a destabilization of the ground state that in turn increases the Boltzmann occupancy of the exchange-competent transiently unfolded states [38]. It is, however, beyond the scope of the present work to elucidate the exact mechanism behind this intriguing allosteric effect.

## Discussion

In this study, we have addressed the dynamics of β-clamp and how it is influenced by ligand binding. Central to this, we obtained assignment of the backbone NMR resonances of β-clamp using a strategy including protein deuteration, TROSY-based NMR spectra, GOODCOP/BADCOP and NOESY connections. As a result, we have NMR reporters distributed throughout this large 83 kDa dimeric protein to monitor structural and dynamic properties. Recently, an ILV methyl NMR assignment of β-clamp was published [39], in which methyl ^13^C and ^1^H chemical shifts of Ile, Leu and Val side chains were assigned. The assignment process of our work could be useful for other large, folded proteins. However, a nearly complete assignment was only possible with the addition of a small, eight-residue peptide ligand comprising the C-terminal CBM of pol III α, highlighting that β-clamp in its free state undergoes NMR-unfavourable dynamic processes.

Addition of a small ligand of only eight residues had a stabilizing effect on β-clamp, affecting its global unfolding process in urea and affecting the dynamics of the amides both locally in the binding site and distant to it. This suggests that binding of a ligand imposes an allosteric effect that redirects dynamics from the binding pocket and the associated loops to the dimer interface, where amide exchange rates increased.

According to our data, the three domains of β-clamp display similar dynamics with similar deuterium uptake. This contrasts two studies by Fang et al. [25,26] who concluded that domain I was the more dynamic. Since local HDX-MS data is not residue specific, and because the deuterium uptake is an average over an entire peptide, the interpretation may depend on the peptides chosen to represent the exchange as well as their length. Therefore, as the peptide covering the β-strand in the dimer interface of domain I is also connected to the highly dynamic IDCL between domains I and II, domain I may appear more exposed and dynamic in their studies as the peptides were longer. Here, we recovered peptides that were shorter and covered the binding interface separately from the IDCL, revealing that the dimer interface is indeed very protected compared to the rest of the protein. Although measured at different conditions (salt and temperatures), we found other discrepancies between our data and those of the Fang et al. papers, which suggest that there is a difference in stability of the β-clamp used here and the one used in the previous studies by Fang et al. One remarkable difference is the presence of EX1 kinetics in the Fang et al data, suggesting global unfolding in the absence of urea, a scenario we do not observe. A more elaborate discussion on these inconsistencies is provided in the Supplementary Information.

Despite its relatively stable structure, we see that β-clamp tends to precipitate at high concentrations. This self-association was reflected in the *R*_2_-values of unbound β-clamp that increased upon increasing concentration (Figure 3C). The self-association was reduced upon pol III binding. This may be because the pol III peptide either screens the hydrophobic binding pockets or prevents local unfolding that exposes hydrophobic residues that may cause β-clamp to self-interact. However, since this self-interaction occur at high concentrations, the phenomenon may not have any biological implications. By HDX-MS we do see that pol III binding has a stabilizing effect on β-clamp by altering the dynamics of β-clamp around the binding pocket and by imposing an overall stabilizing effect on global unfolding in the presence of denaturant.

Generally, we see different types of dynamic processes occurring at different timescales; we see a change in dynamics at fast timescale, reflected by the peak intensity increase for some residues in the NMR spectra, and we see changes in dynamics and stability at the seconds-to-hours timescale with HDX-MS. Residues Ala147, Tyr153 and Gly157 show substantial peak intensity increases in the pol III-bound state. However, since their peak intensities are too low in the free state, we cannot accurately determine an *R_2_*-value for these residues and thus cannot extract at which timescale this occurs at. Yet, these dynamic processes may relate to each other. For example, the protection of residues within the binding pocket caused by pol III binding results in a long-term stabilization of residues more distal to the binding pocket. This allosteric effect even manifests in changes in the dimer interface, which is relatively distant (∼18 Å) from the binding pocket.

Since this global effect on the chemical shifts was also induced by other CBM-containing peptides, we speculate if this change in dynamics may be a common mechanism for ligand binding to β-clamp. As a hub, β-clamp interacts with many different binding partners with different structures and variations in their CBMs that should ultimately lead to a different biological output. The fact that the CBM length vary between five of six residues may further require some structural plasticity. We therefore reason that the flexibility and dynamics of β-clamp is a way of accompanying and adapting to its many different binding partners, and that different binding partner may impose different allosteric effects. The differences in chemical shifts induced by the pol III* and pol IV* peptides may reflect such variable structural adaptability upon binding these different ligands. Structural plasticity and ligand adaptability have also been reported in hub proteins that function as coregulators of sets of transcription factors [40,41]. We also found that oxidation of two methionines in the binding pocket affects the affinity of peptide ligands. One can therefore speculate that these methionine oxidations can have a biological relevance e.g., in partner selection under oxidative stress conditions, but further experiments are needed to investigate if methionine oxidation has a regulatory function in β-clamp interactions.

The β-clamp binding pocket is constituted of many loops and flexible linkers which connect different parts of the structure originating from both domain II and III (Figure 9). Such intricate architecture equips the structure with different structural modulators that can effectuate the information of ligand binding into changes in dynamics at specific sites far away from the binding pocket. Thus, dynamic allostery [42–44] may allow for ligand-dependent modulation of the binding pocket and controlled dissemination of information throughout the protein. This has wide implications and could suggest a relevant and important role also for the non-conserved residues of the SLiMs – a hitherto neglected part of SLiMs. Many β-clamp binding partners also bind to other parts of the β-clamp surface [7,11,19]. How these contacts affect the dynamics and flexibility in the other domains and how these are disseminated throughout the entire structure of β-clamp may be important for differential binding effects. Thus, binding in the context of the full-length protein is important in many aspects of binding, including changes in dynamics. If and how a differential ligand-induced allosteric modulation exist in the interaction between β-clamp and its arsenal of binding partner is a highly intriguing question that deserves future investigations.

**Figure 9:**
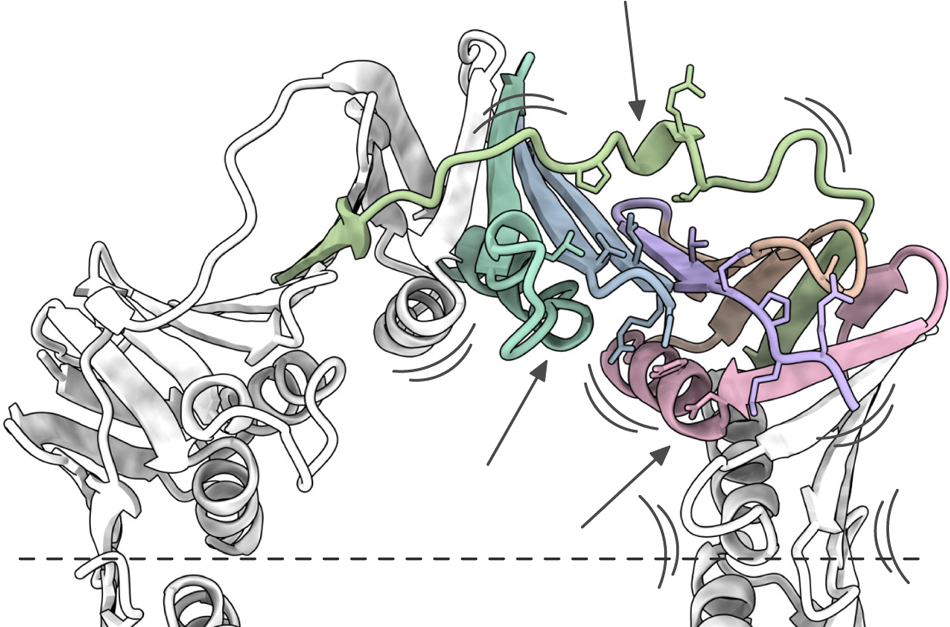
Ligand-induced allosteric effects on β-clamp dynamics. Pol III interacting residues are shown as sticks and different structural segments of the binding pocket is coloured differently. The β-clamp binding pocket is formed by loops of different structured parts which allows for ligand-induced modulation of the dynamics in neighbouring parts of the structure.

DNA sliding clamps are essential proteins in all domains of life [45]. The bacterial β-clamp and the eukaryotic DNA sliding clamp, proliferating cell nuclear antigen (PCNA) are functionally and structurally conserved. PCNA is a trimer, with each subunit comprising of two structurally similar domains [46]. Although all DNA sliding clamps are structurally similar, their intrinsic stability differs [21]. The dynamic properties of PCNA even differ between organisms, where human PCNA is more dynamic, and experiences higher deuterium uptake compared to yeast PCNA; both demonstrated by HDX-MS [26] and HDX NMR [47]. Notably, HDX-MS experiments demonstrated that the binding of a small protein to *Thermococcus kodakarensis* PCNA resulted in increased deuterium uptake in two regions, one of which is a β-strand in the subunit interfaces of the PCNA trimer [48]. Furthermore, NMR studies on human PCNA have revealed substantial effects upon ligand binding even at sites distant to the binding pocket [49–51], comparable to the observations made here for β-clamp interactions. These general observations of allosteric effects indicate that the structural plasticity and allosteric binding mechanism may be common to DNA sliding clamps.

## Materials and methods

### Peptides

The eight-mer pol III peptide (Ac-EQVELEFD) was purchased from Innovagen (Sweden). The other 15-16-mer peptides pol III* (DWRGLIGSEQVELEFD), pol IV* (GWLDPQMERQLVLGL), MutL* (GWGEAPVCAQPLLIPL-NH_2_) and MutS* (GWATQVDGTQMSLLSV-NH_2_) were purchased from TAG Copenhagen (Denmark). The 15-16-mer peptides were designed to contain DW/GW located distant to and N-terminally upstream of the CBM for easier concentration measurements. The 8-mer pol III peptide was N-acetylated to remove the positive charge at the N-terminus, since this is not present in the context of the full-length protein. MutL* and MutS* were C-terminally acetylated since the CBMs of these proteins are internally positioned in the sequence, whereas the CBMs of pol III and pol IV are in the C-terminus of the protein.

### Protein expression and purification

The His_6_-tagged *E. coli* β-clamp sequence in a pET16b vector was transformed into BL21 (DE3) pLysS cells. The full sequence of the N-terminal Hig-tag is (MGHHHHHH). Cells were grown on an LB agar plate containing 100 μg/ml ampicillin (Amp) and 35 μg/ml chloramphenicol (Cam). One colony was added to 10 ml LB media with antibiotics and the culture grown overnight (o/n) at 37°C at 180 rpm. For expression of unlabelled β-clamp, the 10 ml o/n culture was added to 1L LB media supplemented with antibiotics. For expression of ^2^H, ^13^C, ^15^N labelled β-clamp, a pre-culture was made by adding 500 μl LB o/n to 20 ml deuterated M9 media containing 2 g/l ^13^C, ^2^H labelled glucose and 1 g/l ^15^NH_4_Cl in ∼97% D_2_O with 50 μg/ml Amp and 17.5 μg/ml Cam. The pre-culture was grown o/n at 37°C, 180 rpm and added to 0.5 L deuterated ^2^H, ^13^C, ^15^N labelled M9 media the next day. Cells were grown at 37°C, 180 rpm and at OD_600_ ∼0.6-0.8, (∼3 hours for cells in LB media and ∼10 hours for cells in deuterated M9 media), expression was induced by the addition of 0.5 mM IPTG. After 5-6 hours of expression, cells were harvested by centrifugation at 5000 g for 15 min and resuspended in 30 ml Buffer A (20 mM sodium phosphate, 1 M NaCl, 50 mM imidazole, 5 mM β-mercaptoethanol, pH 7.4) including a complete EDTA free protease inhibitor cocktail tablet (Roche Diagnostics GmbH). Cells were lysed by sonication (80% amplitude, 1 cycle for 10x 15 sec with 15 sec breaks between). The lysate was cleared by centrifugation at 20.000 g for 30 min and the lysate transferred onto a gravity column containing 5 ml Ni^2+^-NTA resin (Cytiva Sweden AB) equilibrated with Buffer A. The column was washed with 50 ml Buffer A and the protein eluted in 10 ml Buffer B (20 mM sodium phosphate, 1 M NaCl, 500 mM imidazole, 5 mM β-mercaptoethanol, pH 7.4). The eluted protein was concentrated with a 15 ml 10000 MWCO spin filter (Merck Millipore Ltd.). The sample was centrifuged at 20.000 g for 15 min to remove precipitated protein prior to being loaded onto a HiLoad 16/600 Superdex 200 size exclusion chromatography (SEC) column pre-equilibrated with 20 mM sodium phosphate, 100 mM NaCl, 1 mM dithiothreitol (DTT), pH 7.4 and the sample eluted into the same buffer. Fractions with β-clamp were assessed with SDS-PAGE and pure fractions pooled and stored at 4°C. β-clamp concentrations were calculated by measuring the absorbance at 280 nm on a NanoDrop ND-1000 spectrophotometer (Thermo Fisher Scientific) and using an extinction coefficient of 15930 as calculated by the ExPASy ProtParam web tool (https://web.expasy.org/protparam/).

### NMR spectroscopy

All NMR samples were prepared by adding 5% (v/v) D_2_O and 125 μM DSS to protein solutions in 20 mM sodium phosphate, 100 mM NaCl, 5 mM DTT, pH 7.4. After centrifugation, samples were transferred to 5 mm Shigemi NMR tubes and spectra recorded at 37°C on a Bruker Avance III 750 MHz spectrometer equipped with a cryoprobe. DSS were used to reference ^1^H and ^15^N and ^13^C was referenced indirectly using the gyromagnetic radii. 2D TROSY-^1^H-^15^N-HSQC spectra were processed in Topspin (Bruker). *T*_2_ relaxation experiments and triple-resonance assignment spectra were processed, and phase corrected using qMDD [52] or nmrDraw, a component of NmrPipe [53]. All spectra were analysed and manually assigned in CcpNmr Analysis [54].

Free β-clamp was assigned from a sample containing 1083 μM ^2^H, ^13^C, ^15^N labeled β-clamp using deuterium decoupled TROSY variants of HSQC, HNCA, HNCO as well as ^15^N edited 3D-TROSY-NOESY and gradient optimized CO decoupling pulse (GOODCOP) and beta/alpha decoupling pulse (BADCOP) spectra [31]. A sample of 610 μM ^2^H, ^13^C, ^15^N labelled β-clamp + ∼2-fold excess of pol III was used for the assignment of pol III bound β-clamp using the same experiments as above including a TROSY version of HNCACB. The assignments are deposited to BMRB under the deposition number ID 52494 and 52495. Spectra of β-clamp in complex with pol III*, pol IV*, MutL* and MutS* (**Figure S7**) contained 175 μM ^2^H, ^13^C, ^15^N β-clamp and 700 μM pol III*, pol IV* or MutL* or 350 μM MutS*. MutS* was added to a lower concentration due to low solubility of the peptide.

### Chemical shift perturbations, intensity ratios and secondary chemical shifts

CSPs and peak intensity ratios from pol III binding were quantified from samples of 200 μM ^2^H, ^13^C, ^15^N β-clamp with and without 400 μM pol III. CSPs from addition of pol III* or pol IV* were quantified from titrations into 200 μM ^2^H, ^13^C, ^15^N β-clamp with the addition of up to 800 μM pol III* or pol IV*. Identification of β-clamp peaks in the pol III* and pol IV* bound state were based on a titration with increasing amounts of pol III*/pol IV*. For peaks in intermediate exchange, the assignments were based on the peak trajectory of the first titration point. For the pol III* assignment, peaks in intermediate exchange were also guided by the pol III assignment. Peak intensities were only included for residues where the peak volume could be fit to a parabolic. CSPs were calculated using the following equation [55]:

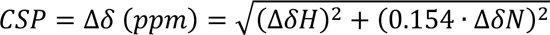

Secondary chemical shifts were calculated using the following equation:

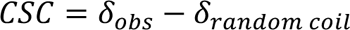

where the intrinsic chemical shifts of a random coil were calculated using the POTENCI web tool (https://st-protein02.chem.au.dk/potenci/) [56] with the physical parameters set to match the conditions of the NMR samples.

### NMR relaxation

A TROSY variant of the *T*_2_ experiment was used to determine *R_2_* relaxation rates of β-clamp with and without pol III. The relaxation delays were 8, 17, 25, 34, 51, 68, 102 and 136 ms and the spectra were recorded in triplicate in a randomized order. The peak heights at each relaxation delay were extracted and fitted to a one phase exponential decay in Prism to obtain the *R_2_* values. Errors are standard error between the three replicas.

### ITC

β-clamp was buffer exchanged into ITC buffer (20 mM sodium phosphate, 100 mM NaCl, 1 mM tris (2-carboxyethyl)phosphine (TCEP), pH 7.4) using a 15 ml 10000 MWCO spin filter (Millipore). The peptides were dissolved in ITC buffer, and the pH measured and adjusted if needed. The cell contained 25.3 – 29.8 μM β-clamp for the titrations with Pol III* and 52.5 – 57.4 μM β-clamp for the titrations with Pol III. The concentration of the pol III* peptide was determined using the extinction coefficient of 5500 as calculated by the Expasy ProtParam web tool (https://web.expasy.org/protparam/) and the measured absorbance at 280 nm from a NanoDrop ND-1000 spectrophotometer (Thermo Fisher Scientific). Protein and peptide samples were centrifuged at 20000 g for 15 min to degas the samples. For all experiments, β-clamp was added to the cell and the peptide was added to the syringe. The ITC experiments were recorded on a Malvern MicroCal PEAQ-ITC instrument (Malvern, United Kingdom) at 25°C and a stir speed of 750 rpm. ITC experiments were repeated 3 times for each peptide.

The MicroCal PEAQ-ITC Software was used to fit the data to a one-site binding model. Because the 8-mer pol III peptide does not contain any aromatic residues, determination of the concentration is more inaccurate and the N-value between the pol III titration and the pol III* titration therefore varied even though the β-clamp stock was identical. Since the measured enthalpy is proportional to the concentration of ligand in the syringe, the N-value was fixed to 1 and the syringe concentration varied to be able to compare the enthalpies between the two peptides.

### HDX-MS

Protein stocks containing 50 μM β-clamp in absence or presence of 500 μM pol III 8-mer peptide in 20 mM sodium phosphate, 100 mM NaCl, 1 mM DTT, pH 7.4 were used for all HDX-MS experiments. Triplicate values were measured for all samples and controls. All hydrogen-deuterium exchange reactions were quenched in formic acid and immediately snap frozen in liquid nitrogen. Samples were kept frozen at −70°C or in liquid nitrogen until loaded onto the LC-MS. For all experiments, each time point represents 50 pmol β-clamp (with or without 500 pmol pol III peptide) that is loaded onto a Waters HDX-Manager with a desalting flow of 0.23% aqueous formic acid solution (Solvent A) provided by an Agilent 1260 Infinity quaternary pump (Agilent Technologies, CA, USA) and a gradient flow comprised of Solvent A (0.23% aqueous FA solution) and Solvent B (0.23% FA in pure acetonitrile, ACN). The gradient was delivered by a nanoAcquity ultra-performance liquid chromatography (UPLC) Binary Solvent Manager (Waters Corporation, MA, USA). Samples were analyzed by electrospray ionization (ESI) MS using an ESI Tri-Wave Ion Mobility mass spectrometer (Synapt G2).

For global and local HDX-MS experiments, non-deuterated controls were made by diluting the β-clamp stock 10-fold in PBS buffer (10 mM phosphate buffer, 2.7 mM KCl, 137 mM NaCl, pH 7.4) and mixing it 1:12 (v/v) with quench solution (0.67% formic acid). Maximally labelled controls were made by diluting β-clamp stock 10-fold in 6M deuterated GdnDCl dissolved in D_2_O for 2 hours at 37°C.

### Global HDX-MS analysis

Protein stocks were diluted 10-fold in deuterated PBS buffer (10 mM phosphate buffer, 2.7 mM KCl, 137 mM NaCl, pD 7.4) and the samples were incubated at 37°C. At 10s, 90s, 810s and 7290s, 10 ul sample was withdrawn and mixed 1:12 (v/v) with quench solution and snap frozen in liquid nitrogen. The thawed samples were afterwards injected into the sample loop (200 μL), desalted for 2 min at 500 μL/min flow rate on a 2.1 mm × 5mm MassPREP Micro Desalting Column at 0.2 °C, and eluted with the following gradient: 5–50% B for 2.5 min, 50–90% B for 0.5 min, 90–95% B for 0.1 min, 95-90% B for 0.3 min and 90-5% B for 0.45 min at a flow rate of 50 μL/min. MS data was acquired for 7 min in positive resolution mode at m/z 300-2500 with a scan time of 0,5 sec. Combined spectra were processed in MassLynx using MaxEnt1 with a width at half height set to 0.45 (0.3 for H_2_O controls). The most populated mass of the non-oxidized state was plotted.

### Urea HDX-MS experiments

Protein stocks were diluted 10 times in deuterated 4M urea, and the samples were incubated at 25°C to avoid covalent adducts. At 10s, 1h, 2h, 4h and 24h 10 ul sample was withdrawn and mixed 1:12 (v/v) with quench solution and snap frozen in liquid nitrogen and analysed as described for the global HDX-MS analysis.

A pulse labelling experiment was made to test if the unfolding by urea was reversible. Here, protein stocks were diluted 10 times in 4M H_2_O urea and incubated at 25°C for 2 hours. After two hours, the sample was diluted 2-fold in 4M deuterated urea and incubated for 1 minute at 25°C before the reaction was quenched.

The data was processed with MaxEnt1 in MassLynx as described for the global HDX-MS analysis. The population of the partially deuterated and the fully deuterated was determined by smoothing the spectra (Smooth window (channels) +/-: 8; Number of smooths: 2; Smoothing method: Mean), subtracting the baseline (Polynomial order: 15; Below curve (%): 30; Tolerance: 0.01) and creating a centred spectrum (Min peak width as half height (channels): 10; Center method: Top; Create centred spectrum; Centred spectrum: Areas). The relative intensities of the centred spectra were extracted and the percent of the fully unfolded calculated as:

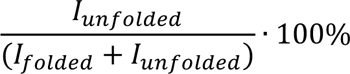

### Local HDX-MS analysis

Protein stocks were diluted 10-fold in deuterated PBS buffer, incubated at 37°C and the reaction quenched after 10s and 7290s in the same way as described above. The samples were injected directly into the sample loop (200 μL) and digested online by agarose immobilized pepsin (ThermoFisher, MA, USA) self-packed in a 2.0 × 20 mm column (IDEX Upchurch guard column, WA, USA) at 20°C. The sample was desalted for 3 min at 300 μL/min flow rate on a ACQUITY UPLC BEH C18 VanGuard Precolumn (130 Å, 1.7 μm, 2.1 × 5 mm, Waters) and separated using 1.0 × 50 mm ACQUITY UPLC Peptide BEH C18 Column (300 Å, 1.7 μm, 1.0 × 50 mm, Waters) at 0.2 °C. The proteolytic peptides were eluted with 5–10% B for 0.1 min, 10–50% B for 11.9 min, 50– 90% B for 0.5 min, 90%-90% B for 0.1 min and 90-5% B for 0.1 min at a constant 40 μL/min flow rate.

Peptide identifications by LC-MS/MS analysis was conducted using an Orbitrap Eclipse mass spectrometer operating in positive ion mode (Thermo FisherScientific, San Jose, CA, USA). The peptides were chromatographically separated using the same equipment and methods as for the local HDX-MS analysis described above. Mass spectra were acquired with an automatic data-dependent switch between an Orbitrap survey MS scan in the mass range of m/z 300 to 1500, followed by fragmentation of peptides using collisional energy of 30% (normalized) in a 1 s cycle. MS1 spectra were acquired at a resolution of 120,000 at m/z 200. A dynamic exclusion of previously selected ions for 5 s was applied. MS2 spectra were obtained at 60,000 resolution at m/z 200 with a normalized AGC target of 300% using a m/z 1.2 isolation window. The maximum injection time was 246 ms (MS1) and 118 ms (MS2). The MS1 AGC target was 400,000 ions, and the MS2 AGC target was 150,000 ions. The resulting raw files were converted to MGF files using ProteoWizard, and the peptides were identified by a Mascot database search with the following parameters: variable modifications (oxidation, M), (Sulfo, STY); peptide mass tolerance ±0.1 Da; fragment ion mass tolerance ± 7 ppm; enzyme, none. The Mascot database contained the proteins of interest as well as possible contaminating proteins.

The local HDX-MS data was analysed in DynamX. Relative deuterium uptakes were extracted from DynamX and plotted into Excel for calculations of uptake and back-exchange. % deuteration was calculated as:

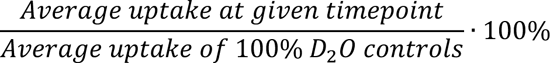

Percent back-exchange was calculated as:

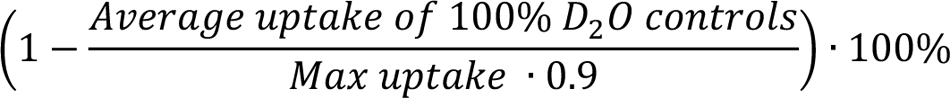

The fraction of bound protein was calculated using the following equation [57]:

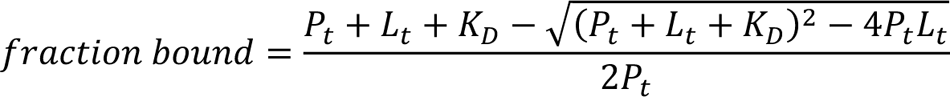

Where *P_t_* is the total protein concentration and *L_t_* is the total ligand concentration.

## Data availability

The chemical shifts of free and pol-II bound E. coli of β-clamp are deposited to BMRB under the deposition number IDs 52494 and 52495 (released upon publication)

## CRediT authorship contribution statement

Signe Simonsen: Conceptualization, Investigation, Formal analysis, Writing – Original Draft, Writing – Review & Editing, Visualization. Andreas Prestel: Conceptualization, Formel analysis, Writing – Review & Editing. Eva C. Østerlund: Investigation, Formel analysis, Writing – Review & Editing. Marit Otterlei: Writing – Review & Editing, Funding acquisition. Thomas J.D. Jørgensen: Conceptualization, Writing – Original Draft, Writing – Review & Editing, Supervision. Birthe B. Kragelund: Conceptualization, Writing – Original Draft, Writing – Review & Editing, Supervision.

## Acknowledgements

We would like to thank Signe A. Sjørup for technical assistance and Johan G. Olsen for discussion. This work was supported by grants from the Novo Nordisk Foundation Challenge program to REPIN (#NNF18OC0033926 to B.B.K.), a generous grant from the VILLUM Foundation to the VILLUM Center for Bioanalytical Sciences at the University of Southern Denmark (to T.J.D.J.) and from the Trond Mohn Research Foundation (to M.O.). All NMR data was recorded at cOpenNMR, Department of Biology, UCPH, an infrastructure supported by the Novo Nordisk Foundation (NNF18OC0032996).

## Abbreviations

NMR: nuclear magnetic resonance

HDX-MS: hydrogen-deuterium exchange mass spectrometry

CBM: clamp binding motif

SLiM: short linear motif

pol: polymerase

GOODCOP: gradient optimized CO decoupling pulse

BADCOP: beta/alpha decoupling pulse

CSP: Chemical shift perturbation

SCS: secondary chemical shift

## Supplementary Information

### Simonsen et al

**Figure S1:**
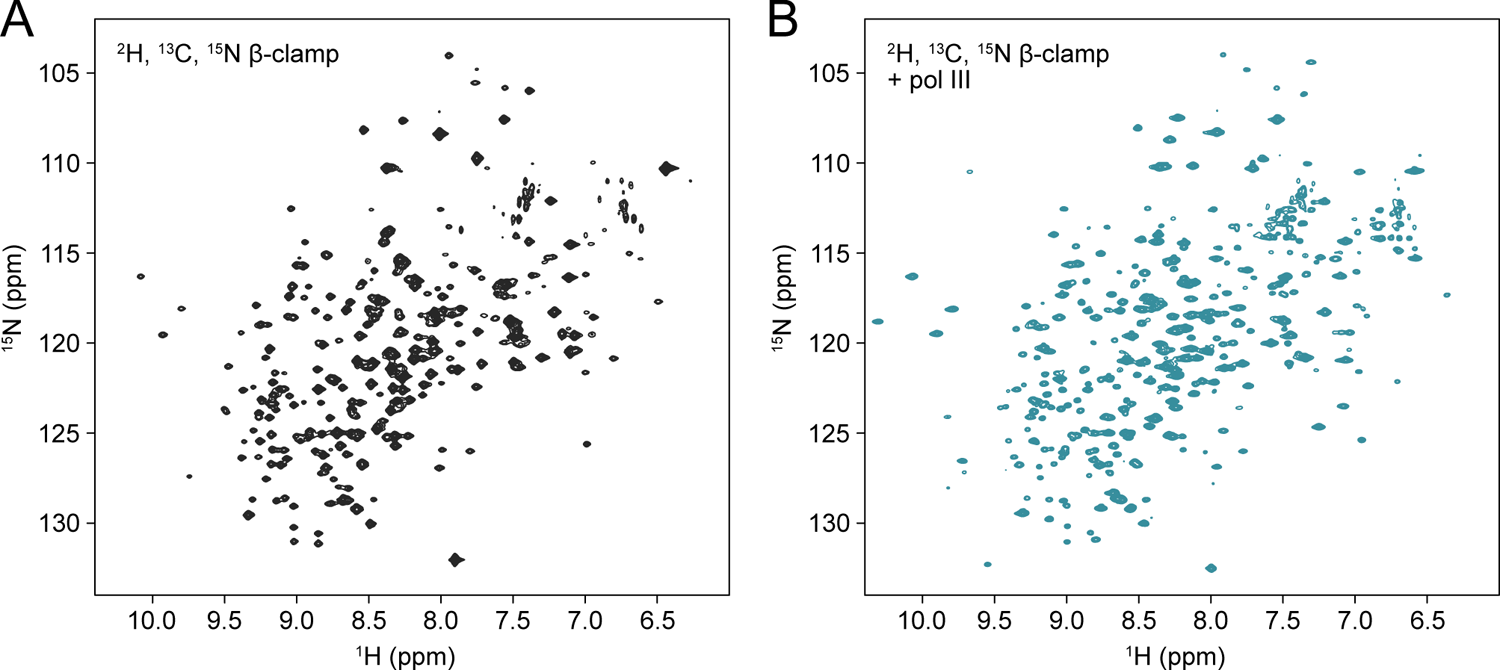
^1^H, ^15^N TROSY-HSQC of ^2^H, ^13^C, ^15^N β-clamp without (A) and with the addition of pol III peptide (B).

**Figure S2:**
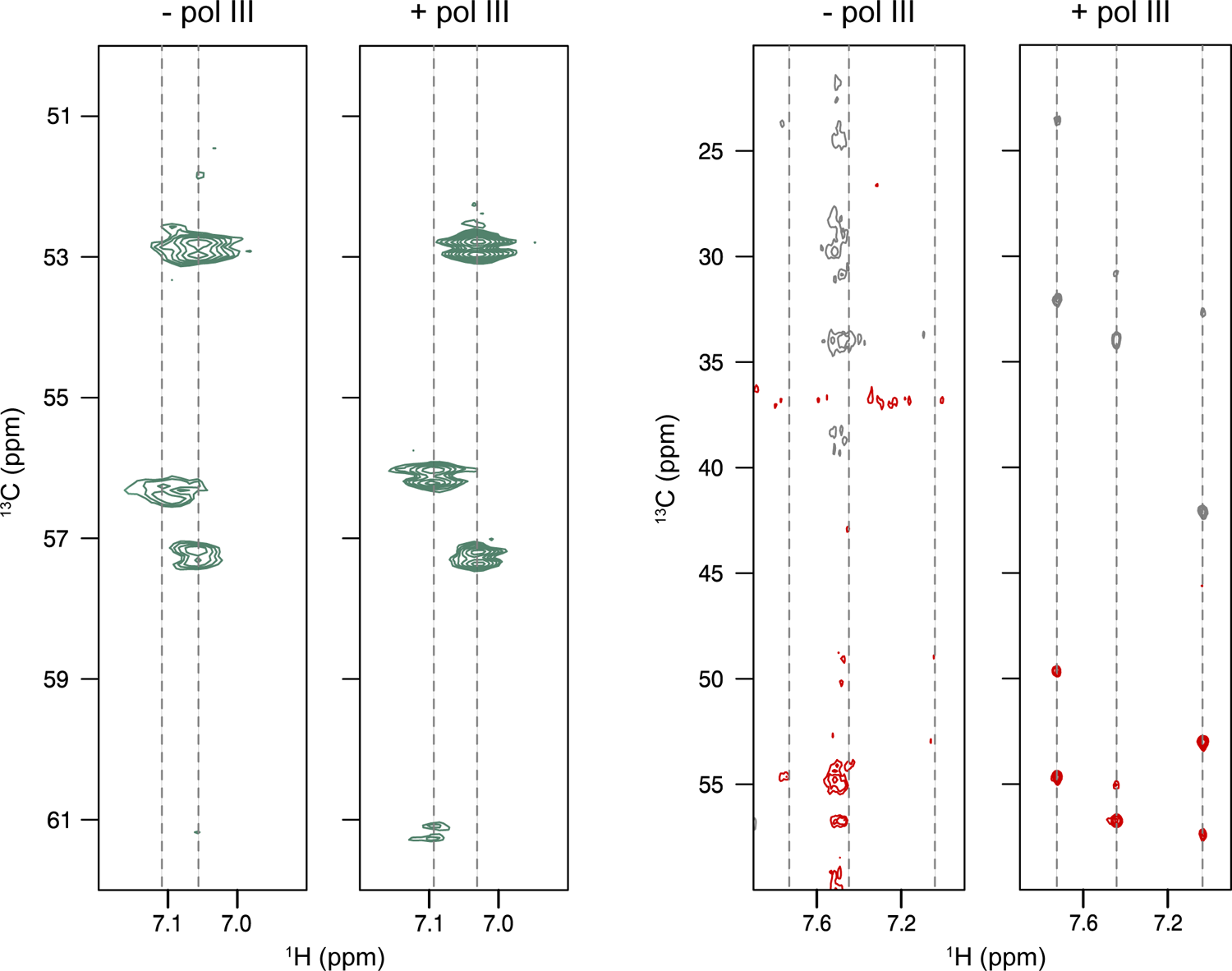
Addition of pol III peptide improved the spectral quality of the 3D assignment spectra. Left: GOODCOP HNCA spectra with and without the addition of pol III. Two peaks are supposed to appear at each line, but a minor peak is missing in the spectrum with free β-clamp. Right: HNCACB spectra with and without pol III. Four peaks, two C^α^ (red) and two C^β^ (grey), are supposed to appear on each line, but the free β-clamp spectrum is pure noise.

**Figure S3:**
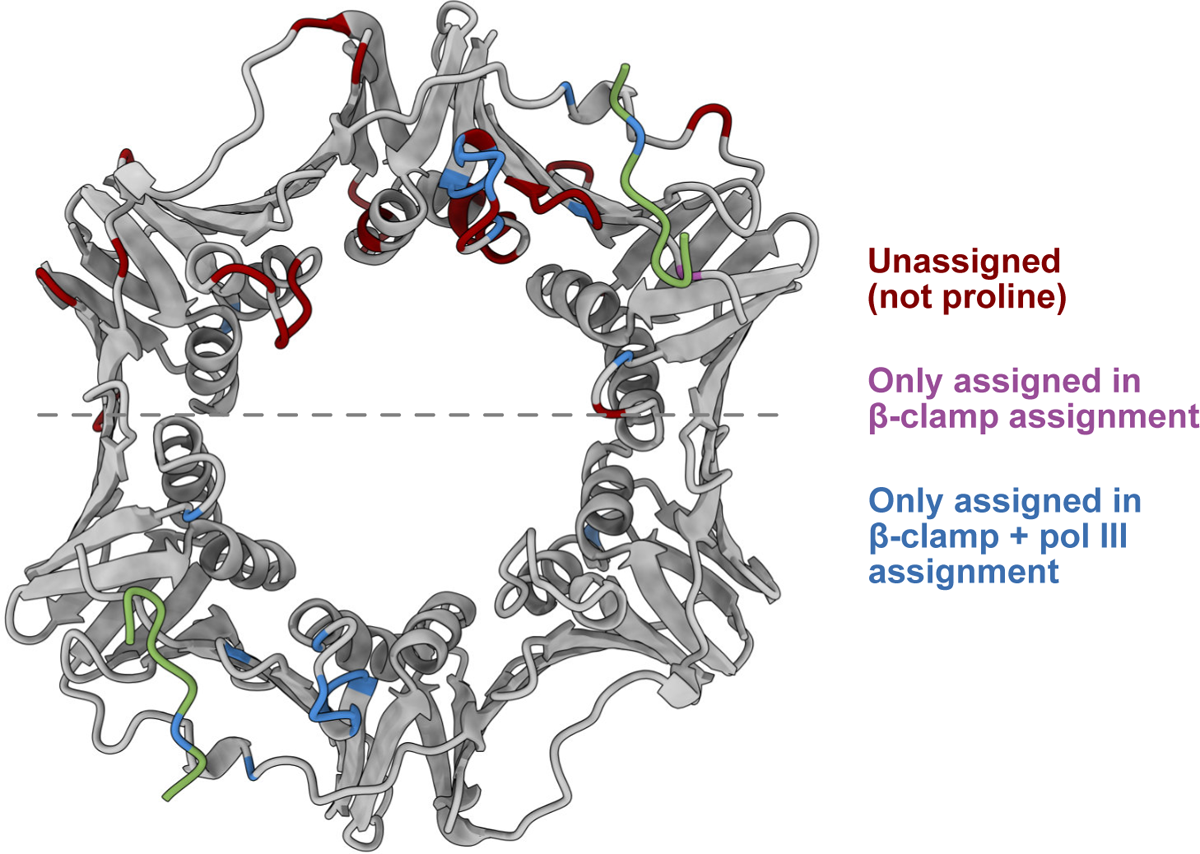
Unassigned β-clamp residues. β-clamp in complex with pol III peptide (green) (PDB ID 3D1F [30]). Unassigned residues are coloured red (not prolines). Residues only assigned in the assignment of pol III bound β-clamp (Glu8, Ala147, Val151, Arg152, Tyr153, Leu155, Asn156, Gly157, Arg176, Asp243 and Phe278) are coloured blue. Met364 which is only assigned in the assignment of unbound β-clamp is coloured purple. Only residues on one β-clamp monomer are coloured.

**Figure S4:**
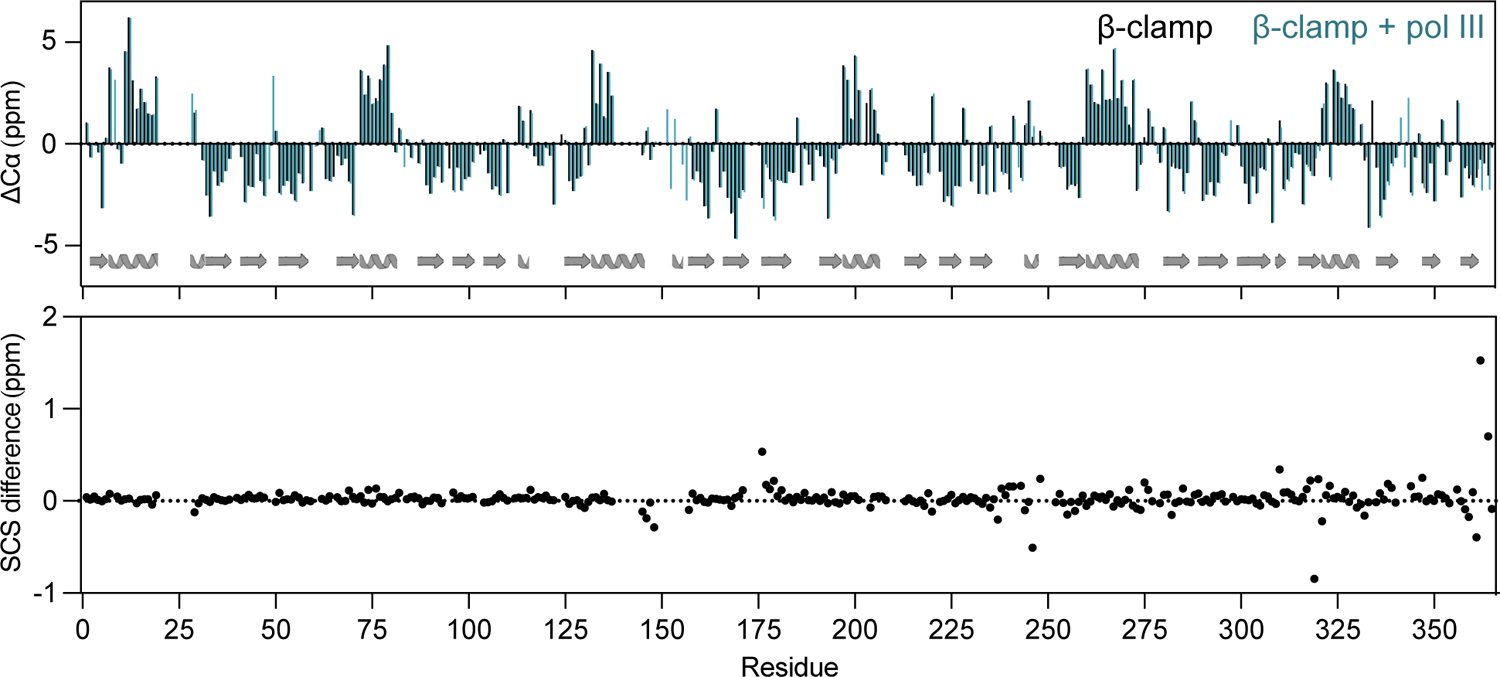
Ligand binding causes no drastic change in secondary structure of β-clamp. Top: Secondary chemical shifts of unbound β-clamp (black) and β-clamp + pol III (blue). Secondary structures are mapped on the sequence according to the β-clamp crystal structure. Bottom: Difference in secondary chemical shifts between unbound and bound β-clamp (SCS_β-clamp_ - SCS_β-clamp + pol III_).

**Figure S5:**
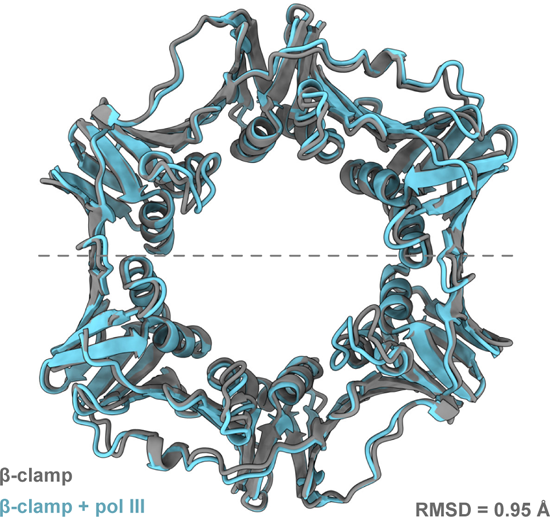
pol III peptide binding does not cause large structural changes in crystal structures of β-clamp. Structural alignment of unbound β-clamp (PDB ID: 1MMI [14]) and β-clamp in complex with pol III peptide (PDB ID: 3D1F [30]).

**Figure S6:**
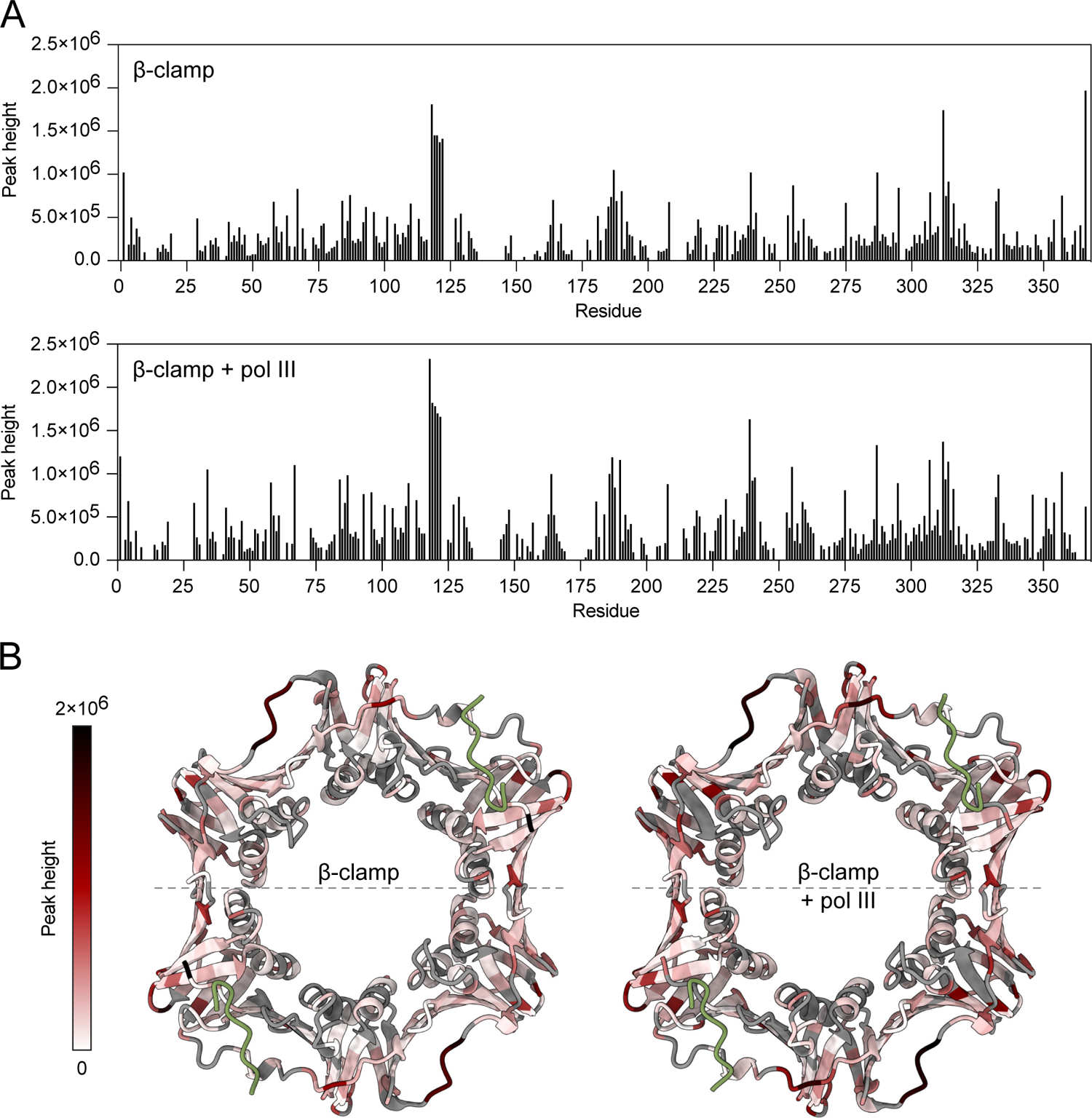
Peak intensities of β-clamp with and without the addition of pol III peptide. A: Peak heights from ^1^H, ^15^N TROSY-HSQCs of ^2^H, ^13^C, ^15^N labelled β-clamp without (top) and with (below) the addition of pol III peptide. B: Peak height mapped onto the structure of β-clamp in complex with pol III peptide (PDB ID: 3D1F [30]). Unassigned residues and residues where peaks could not be fit to a parabolic are coloured grey.

**Figure S7:**
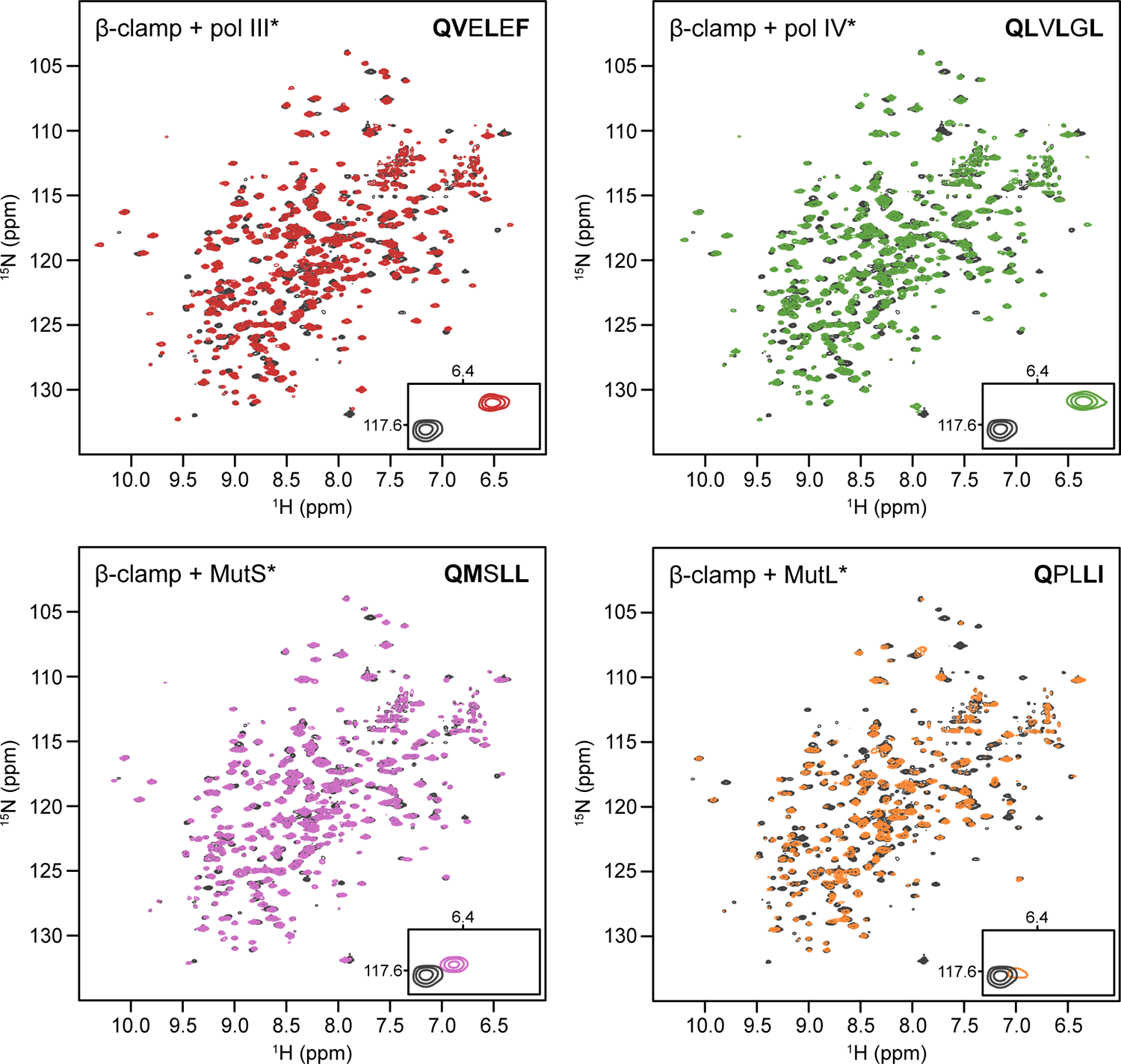
Global CSP effects are common to β-clamp ligands but differ in magnitude. ^1^H, ^15^N TROSY-HSQC of ^2^H, ^13^C, ^15^N labelled β-clamp without (black) and with 2-4 molar excess of peptide ligands (175 μM ^2^H, ^13^C, ^15^N β-clamp + 700 μM pol III*, pol IV*, MutL* or 350 μM MutS*). CBMs of the different binding partners are written in the top right corner with SLiM interacting residues in bold. Bottom right corner shows a zoom of Val335 highlighting the difference in the magnitude of the CSPs and the change in peak intensity in the presence of MutL*.

**Figure S8:**
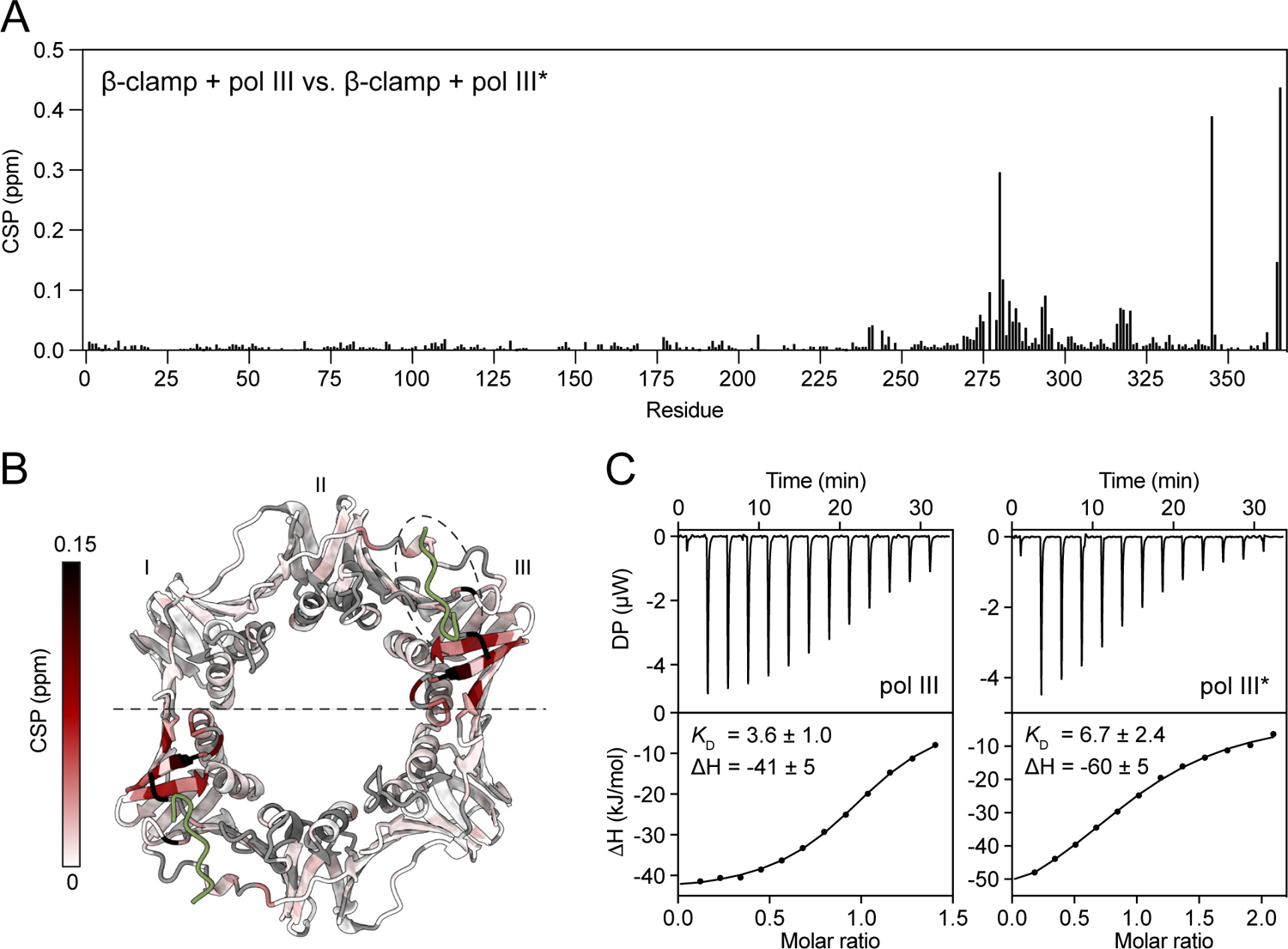
CBM Flanking regions have small effects on binding enthalpy and affinity. A: Chemical shift perturbations from the ^1^H, ^15^N TROSY-HSQCs between ^2^H, ^13^C, ^15^N β-clamp + pol III and ^2^H, ^13^C, ^15^N β-clamp + pol III*. B: Comparison of the chemical shifts in the ^1^H, ^15^N TROSY-HSQCs of β-clamp bound to pol III and pol III*. CSPs mapped onto the structure of β-clamp in complex with pol III peptide (PDB ID: 3D1F [30]) with pol III peptide coloured green and unassigned residues coloured grey. C: ITC data of the titration of pol III and pol III* peptides into β-clamp. The upper parts show baseline corrected raw heats from the titration and lower parts show the integrated binding isotherm and the fitted binding curves. The data was fitted to a ‘one set of sites’-model with the N-value fixed to 1 to allow for comparison between the enthalpies.

**Figure S9:**
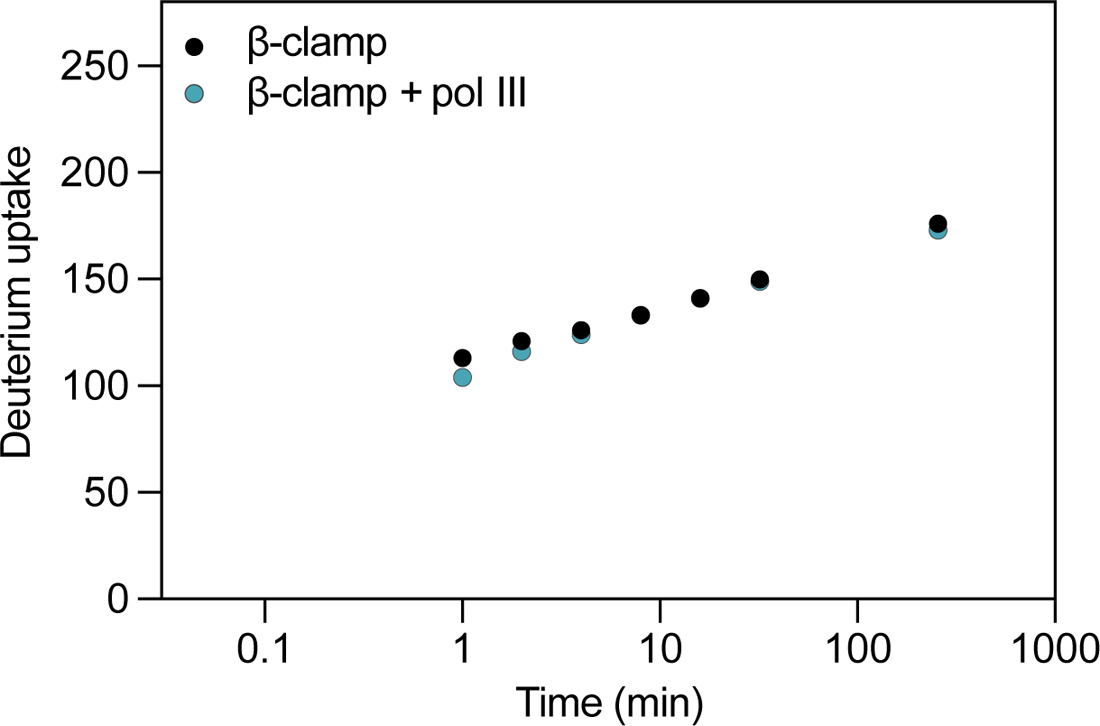
Pilot HDX-MS study on the global exchange of free and pol III bound β-clamp. Deuterium uptake of free β-clamp and pol III bound β-clamp after incubation for 1 min, 2 min, 4 min, 8 min, 16 min, 32 min and 256 min in deuterated PBS buffer at 37°C, pD 7.4. Only one replicate and no maximally labelled sample was made, but the values overall agree with the deuterium uptake in Figure 5. The pilot study was performed under the same conditions as the global analysis in Figure 5, expect that the samples were mixed 1:3 (v/v) with quench solution and that 100 pmol β-clamp was loaded onto the LC-MS.

**Figure S10:**
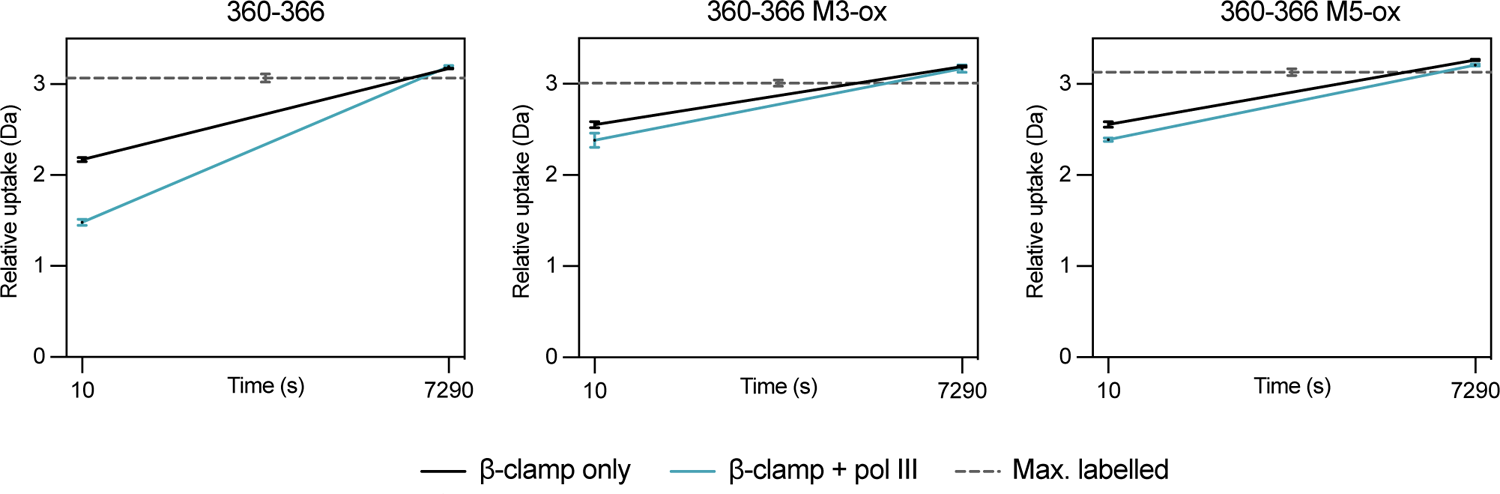
Uptake plots of peptide 360-366 non-oxidized (left), with M362 oxidized (middle) and with M364 oxidized (right). The uptake plots show how the HDX protection after 10 seconds in the presence of pol III is reduced when β-clamp is oxidized in the two methionines in the binding pocket.

**Figure S11:**
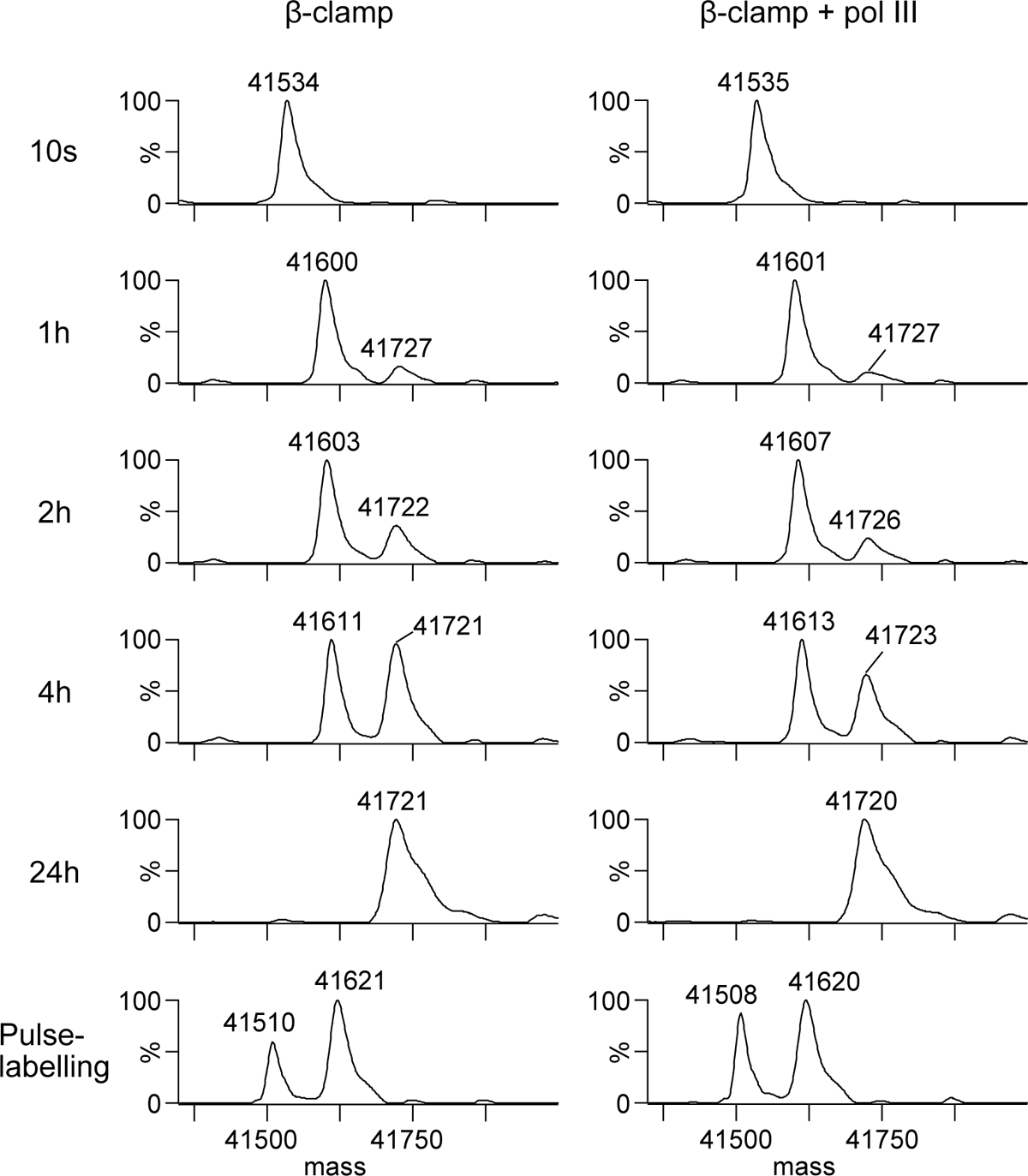
pol III causes slower global unfolding of β-clamp. Deconvoluted ESI-MS spectra obtained from β-clamp incubated in 4M deuterated urea in the presence and absence of pol III. The low-mass population corresponds to folded β-clamp and the high-mass population corresponds to globally unfolded maximally labelled β-clamp. Note that the pulse-labelling experiment shows a different mass, as these samples were only diluted 2-fold in deuterated urea.

**Figure S12:**
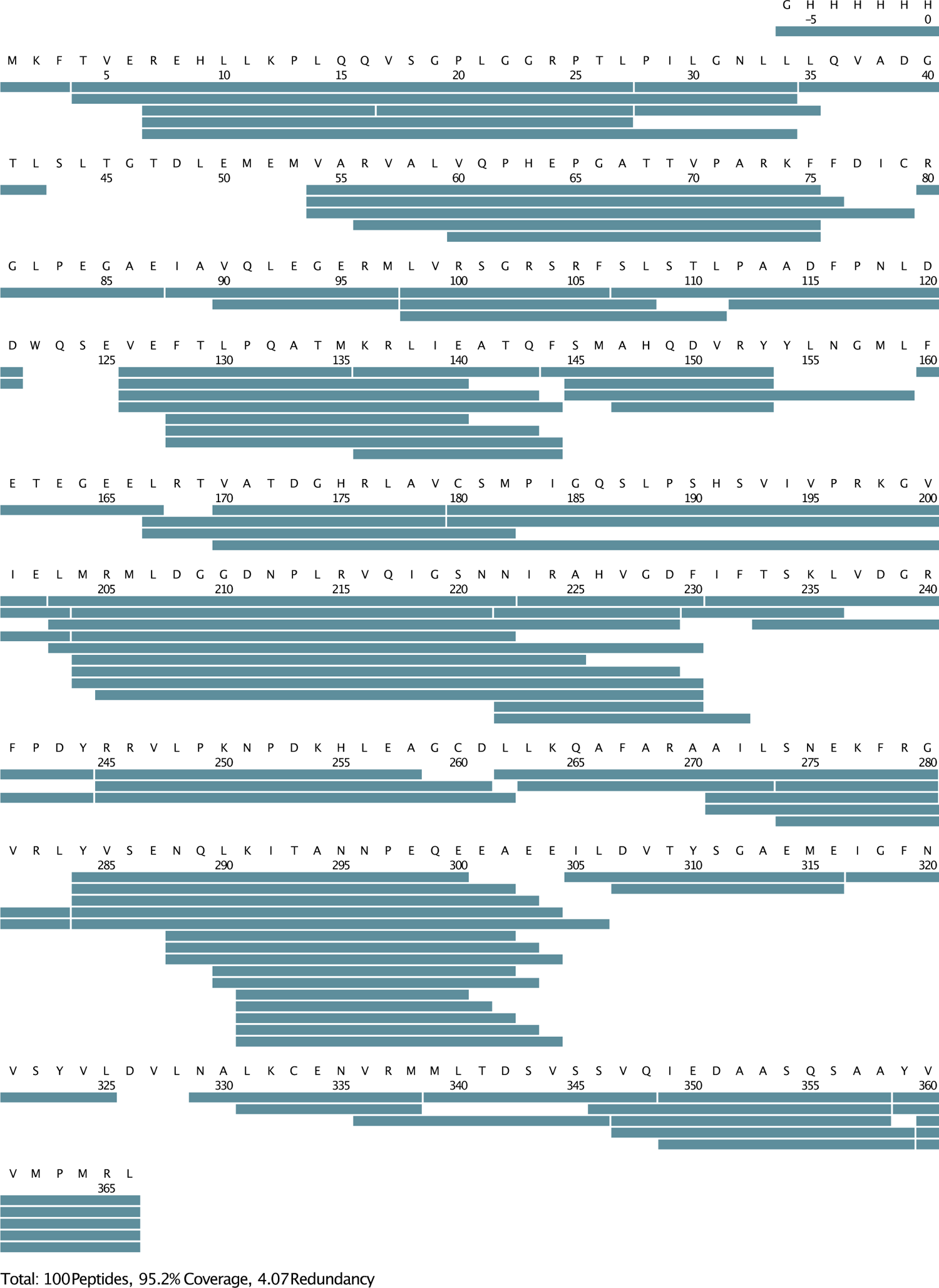
Sequence coverage map of all identified β-clamp peptides for local HDX-MS. All peptides are listed in Table S1. The list includes 100 peptides with 95.2% coverage and 4.07 redundancy. The coverage map was exported from DynamX.

**Figure S13:**
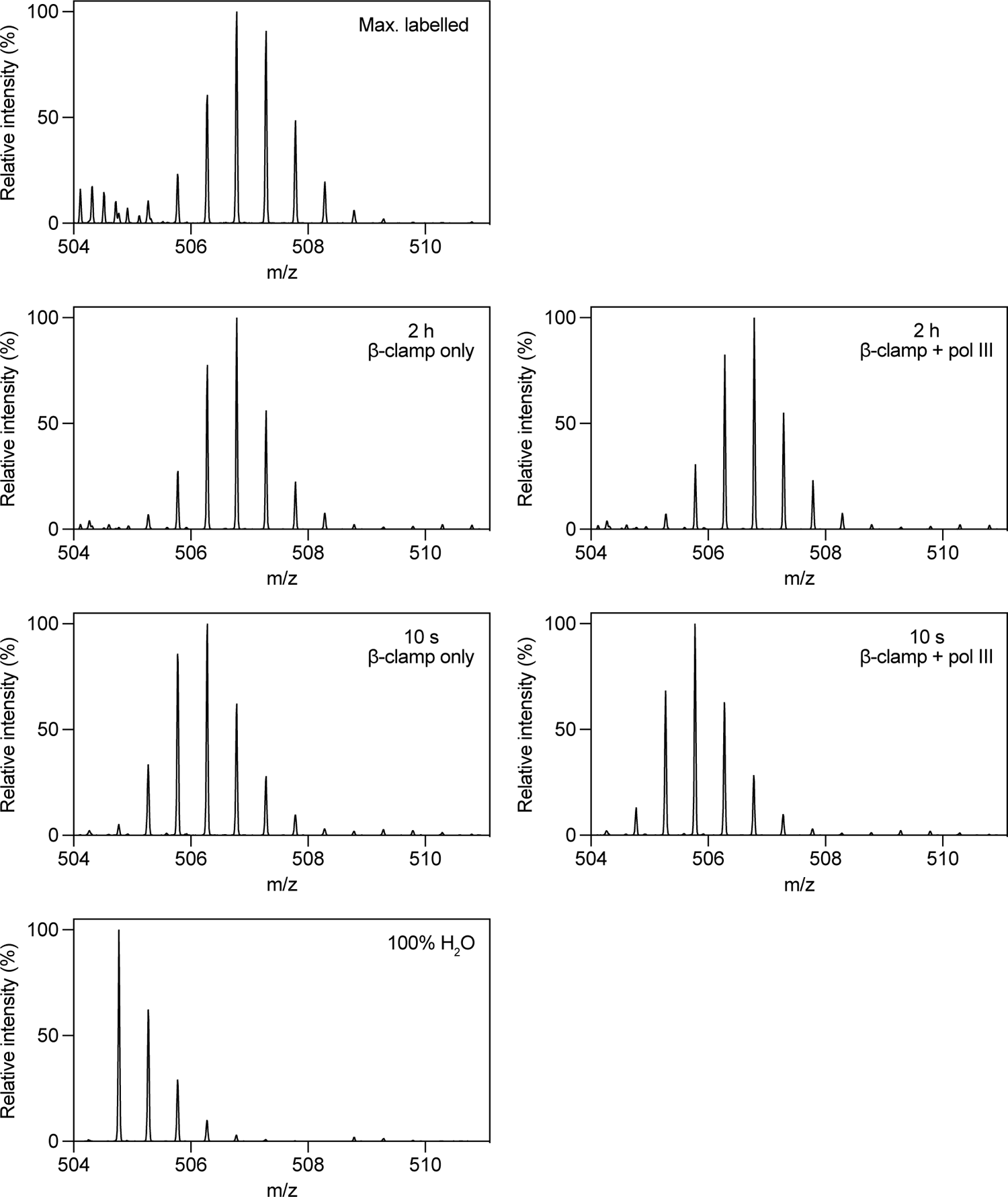
Example of local uptake spectra for peptide 359-366. The plots show the mass/charge of the +2 peptide in the 100% H_2_O control, the maximally labelled control, and β-clamp samples where the HDX reaction is quenched after 10 seconds and 2 hours in the absence (left) and presence (right) of pol III. Plots are exported from DynamX and plotted in Prism.

**Figure S14:**
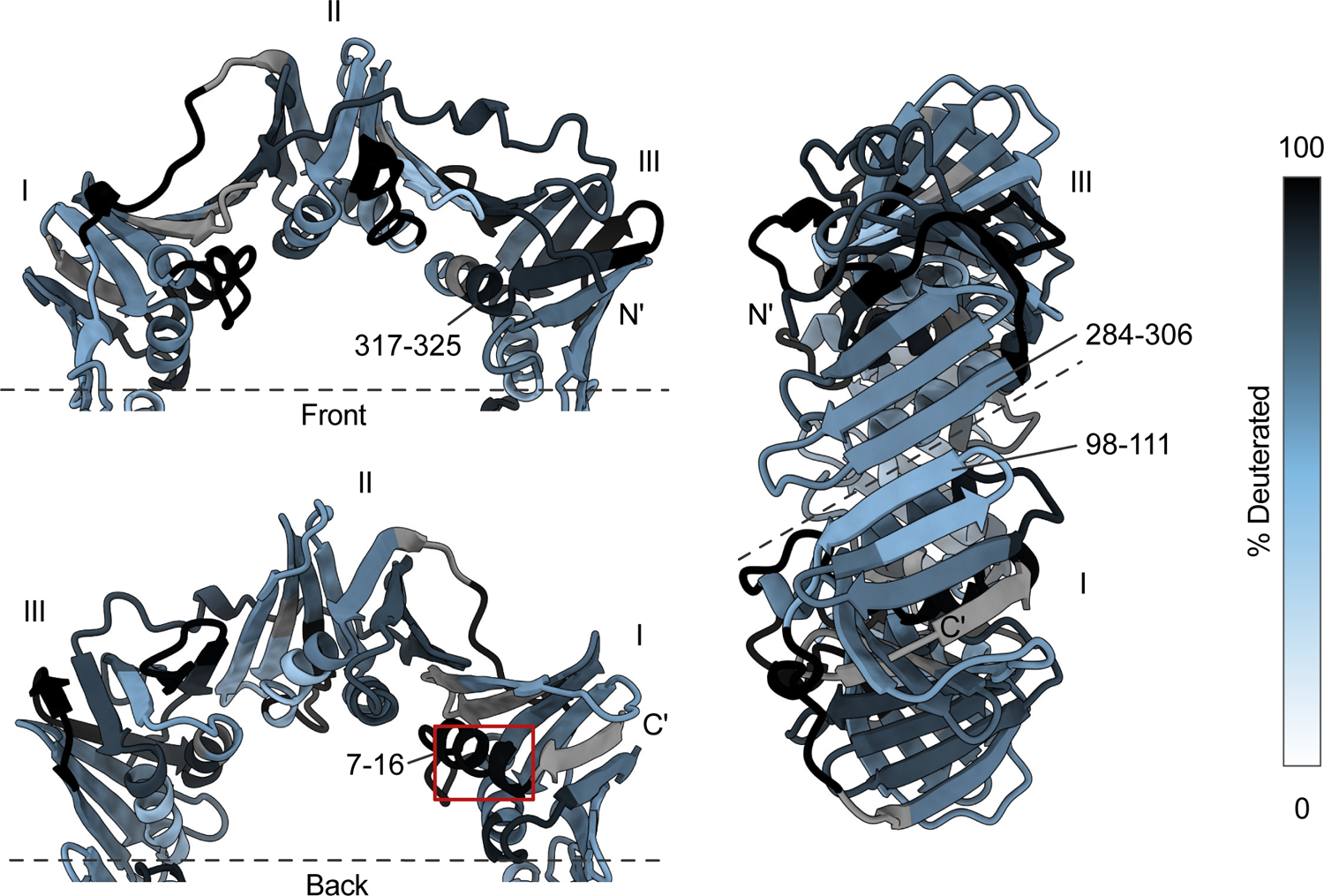
Hydrogen-deuterium exchange mapping with selected peptides highlighted. Peptides 7-16 and 317-325 show high deuterium incorporation despite not containing any loops in the crystal structure. Front side is the site for pol III binding and back side is the opposite site of β-clamp. Peptides 98-111 and 284-306 lie in the in the dimer interface and display relatively low deuterium incorporation compared to the rest of the structure. PDB ID: 3D1F [30]. HDX-exchange level after 2 hours is shown.

**Figure S15:**
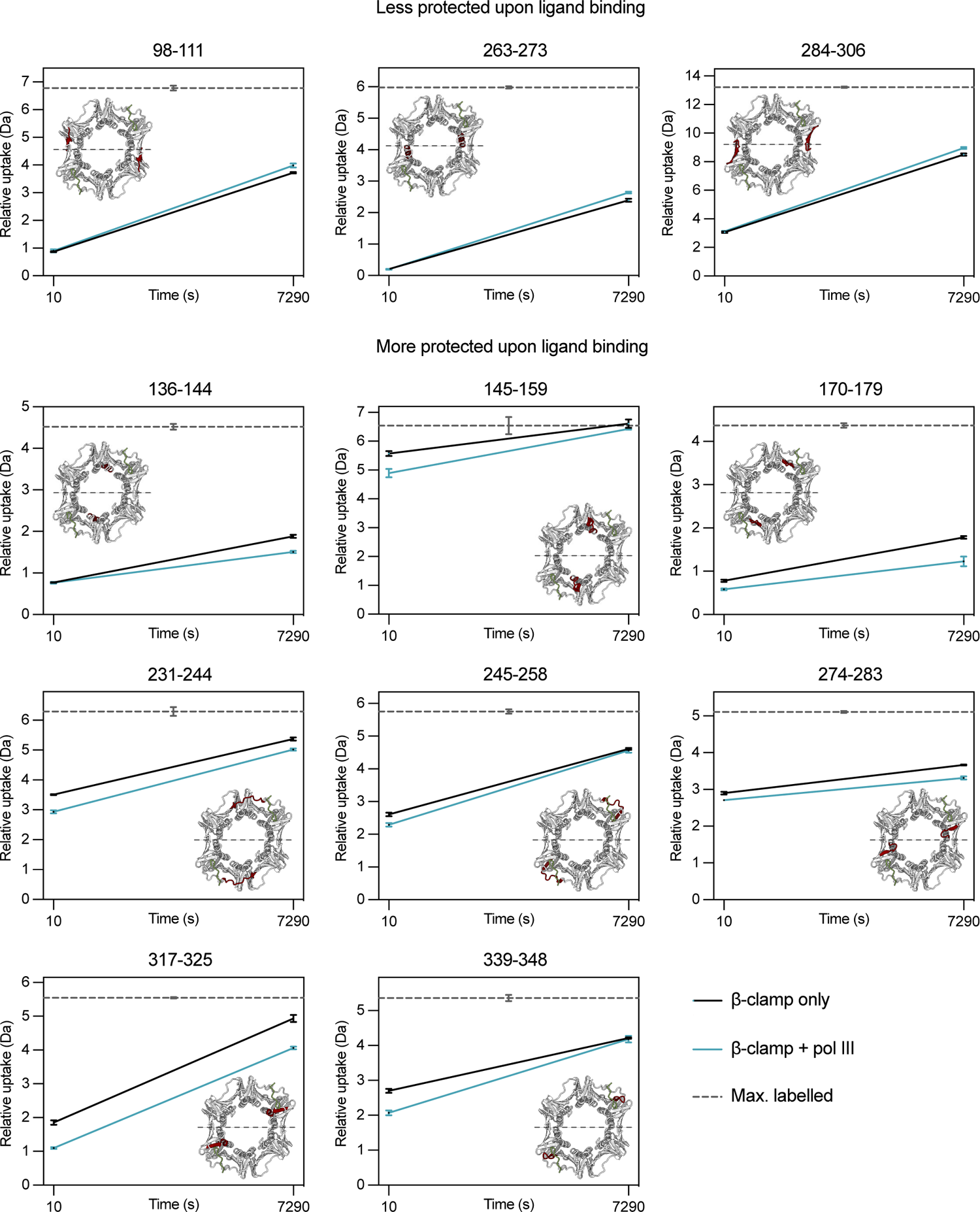
Uptake plots of representative peptides with more deuterium uptake (top row) or less deuterium uptake (three bottom rows) upon pol III binding. The uptake plots show the relative deuterium uptake of β-clamp peptides in absence (black) and presence (blue) of pol III.

### Discussion and comparison with Fang et al. papers

Fang et al [25,26] reported that β-clamp undergoes reversible cooperative unfolding with a half-life of ∼3.5 h at 25°C, pD 7.5 as measured by local HDX-MS analyses. The unfolding of β-clamp yielded bimodal peaks, which is a mass spectral signature of EX1 kinetics. In the present study, we did not observe EX1 kinetics. Because our local HDX-MS analysis only included two time-points and were recorded at 37°C (vs. 25°C in the experiments by Fang et al.) we might be measuring outside the EX1 kinetics regime. However, the reason we don’t see any indication of EX1 kinetics could also likely be caused by a difference in stability between our β-clamp samples and the samples analysed by Fang et al.

Fang et al. estimated a half-life of 3.5 hours for unfolding of the intact β-clamp [25]. Given a half-life of 3.5 hours, ∼60% of β-clamp would have undergone cooperative unfolding by EX1 kinetics after 4 hours at 25°C in a 20 mM HEPES buffer with 50 mM NaCl. In the present study, ∼60% of β-clamp undergoes a cooperatively unfolding after 4 hours at 25°C, pH 7.4 under moderate denaturing conditions, i.e., in PBS buffer with 4M urea (Figure 7). The fact that we observe a similar unfolding rate, but under moderate denaturing conditions, strongly suggests that β-clamp is substantially more kinetically stable in our study than that reported in the study by Fang et al. This increased stability is most likely also the reason as to why we did not observe any indications of EX1 kinetics. In the local HDX-MS analysis of β-clamp at 37°C in physiological deuterated buffer (i.e., without urea), all peaks were unimodal with no apparent peak broadening.

Furthermore, in the present study, the N-terminal part of domain I (peptide 4-27) is completely exchanged after 2h deuteration at 37°C. By contrast, the deuterium uptake of the rest of domain I is significantly below the maximally labelled control. For example, the deuterium uptake of peptide 54-79 is only ∼60% relative to the maximally labelled control after 2h deuteration at 37°C. This reduced deuterium uptake reflects that this part of β-clamp (i.e., residues 28-111) is substantially more protected than the N-terminus (peptide 4-27). In contrast, Fang et al. found that the N-terminus of domain I had similar stability as other parts of domain I and that it exchanged predominantly *via* EX1 kinetics with an overall half-life of 3.5 h at 25°C (as shown by the time-dependent emergence of bimodal peak patterns for, e.g., peptides 1-35, 35-51, 54-89).

Our global analysis on the deuterium uptake is in overall agreement with that of Fang et al. [25]. We see a higher deuterium uptake after 10 s and ∼2 hours, yet this is expected because the higher temperature in our experiments would cause faster exchange. However, Fang et al. either do not see any EX1 kinetics in their global analysis, or they do not comment on it. There are therefore some discrepancies between their global and local HDX-MS data. Furthermore, in the most recent of the two papers, they state that the overall deuterium uptake was higher than in the previous paper, although the EX1 exchange pattern was the same [26]. Their studies also do not include a maximally labelled control, which makes it difficult to estimate the maximal amount of deuteration incorporation, although the exchange of the EX1 globally unfolded β-clamp monomer after six hours serves as an indication for the maximally labelled peptide. A direct comparison of our data and that of Fang et al. is therefore rather complicated.

The underlying reason for the difference in β-clamp dynamics between our study and that of Fang et al. is currently not clear. The ionic strength of the exchange buffer is higher in the present study, but that would be expected to decrease the stability of β-clamp [29]. Additionally, our β-clamp protein contains a N-terminal His_6_-tag. Although we don’t see any indications that the His_6_-tag should form interactions with β-clamp, we cannot completely exclude that it does not have any effect on the intrinsic stability of β-clamp. However, since the His_6_-tag is located far from the pol III binding pocket, we do not expect it to interfere with the interaction with pol III nor affect the allosteric effects we see upon ligand binding.

**Table S1:**
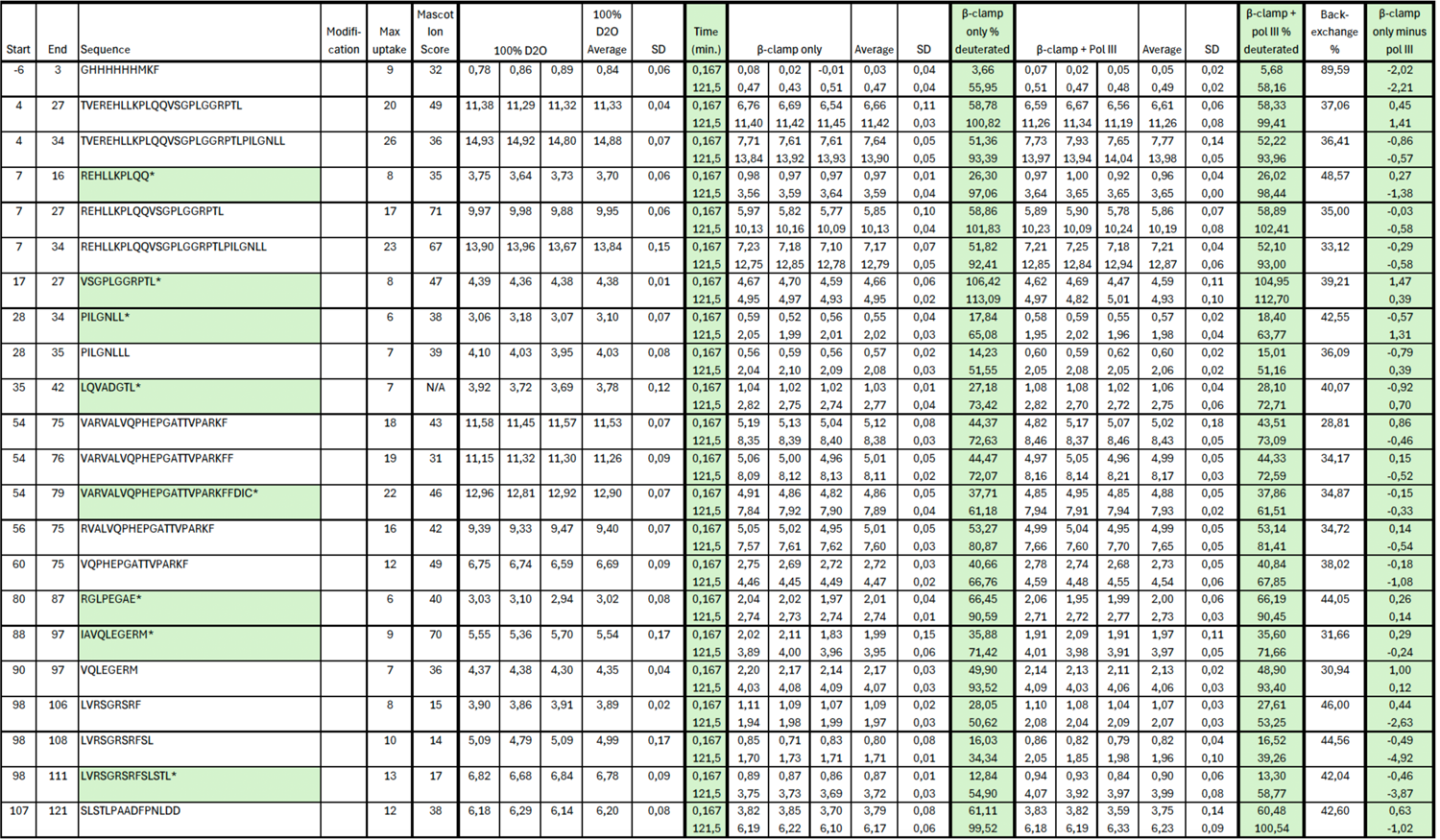
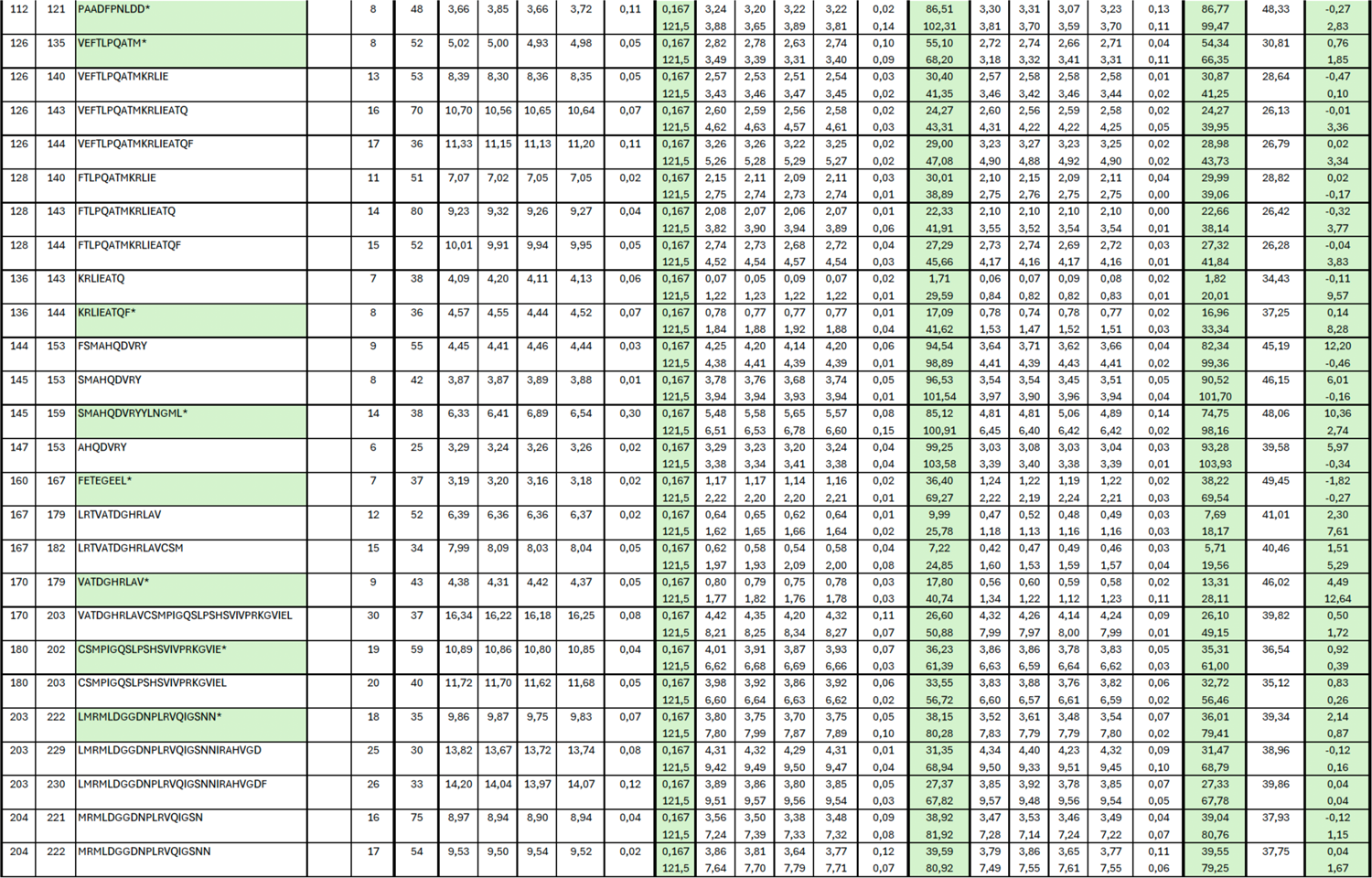
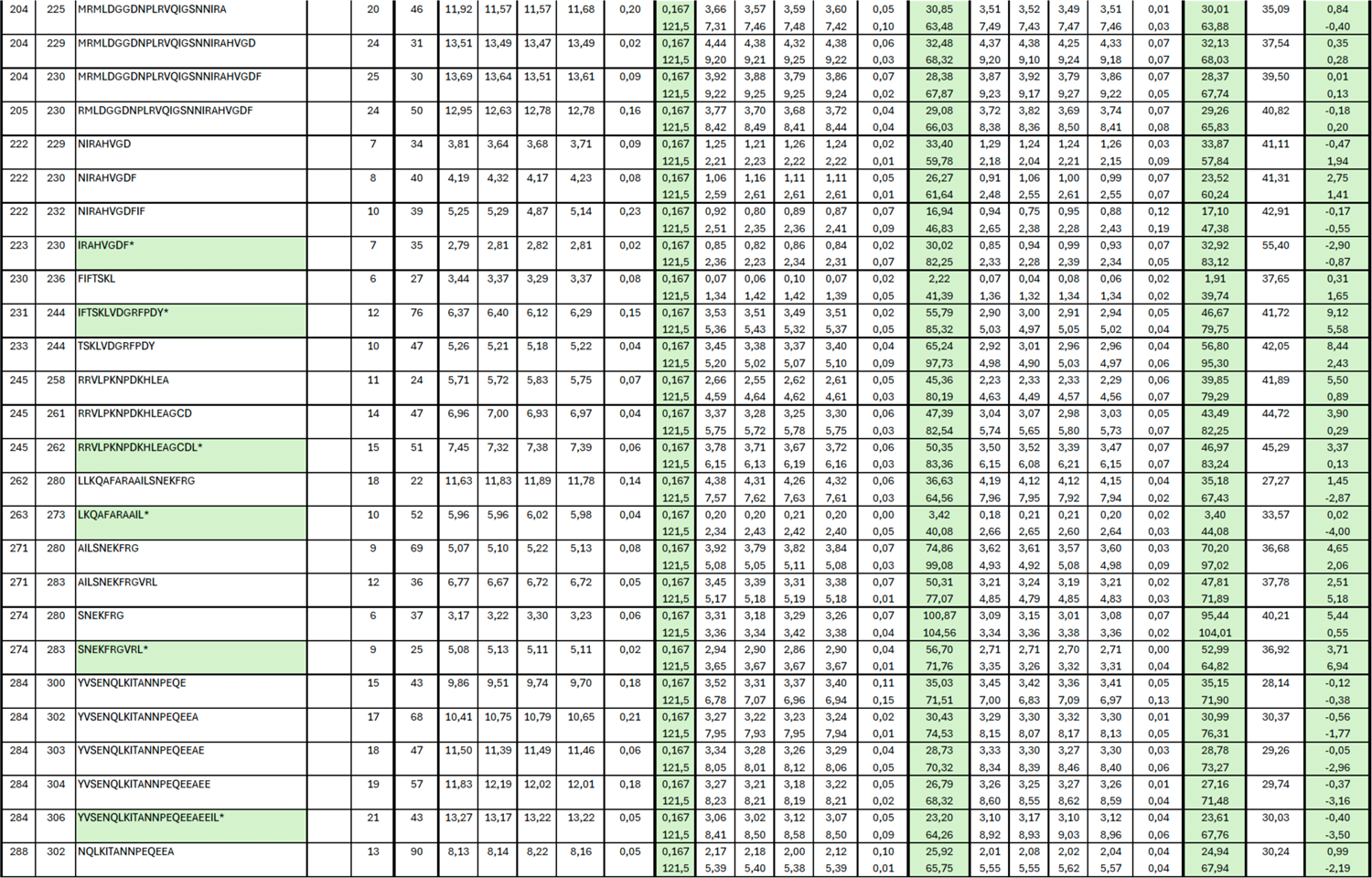
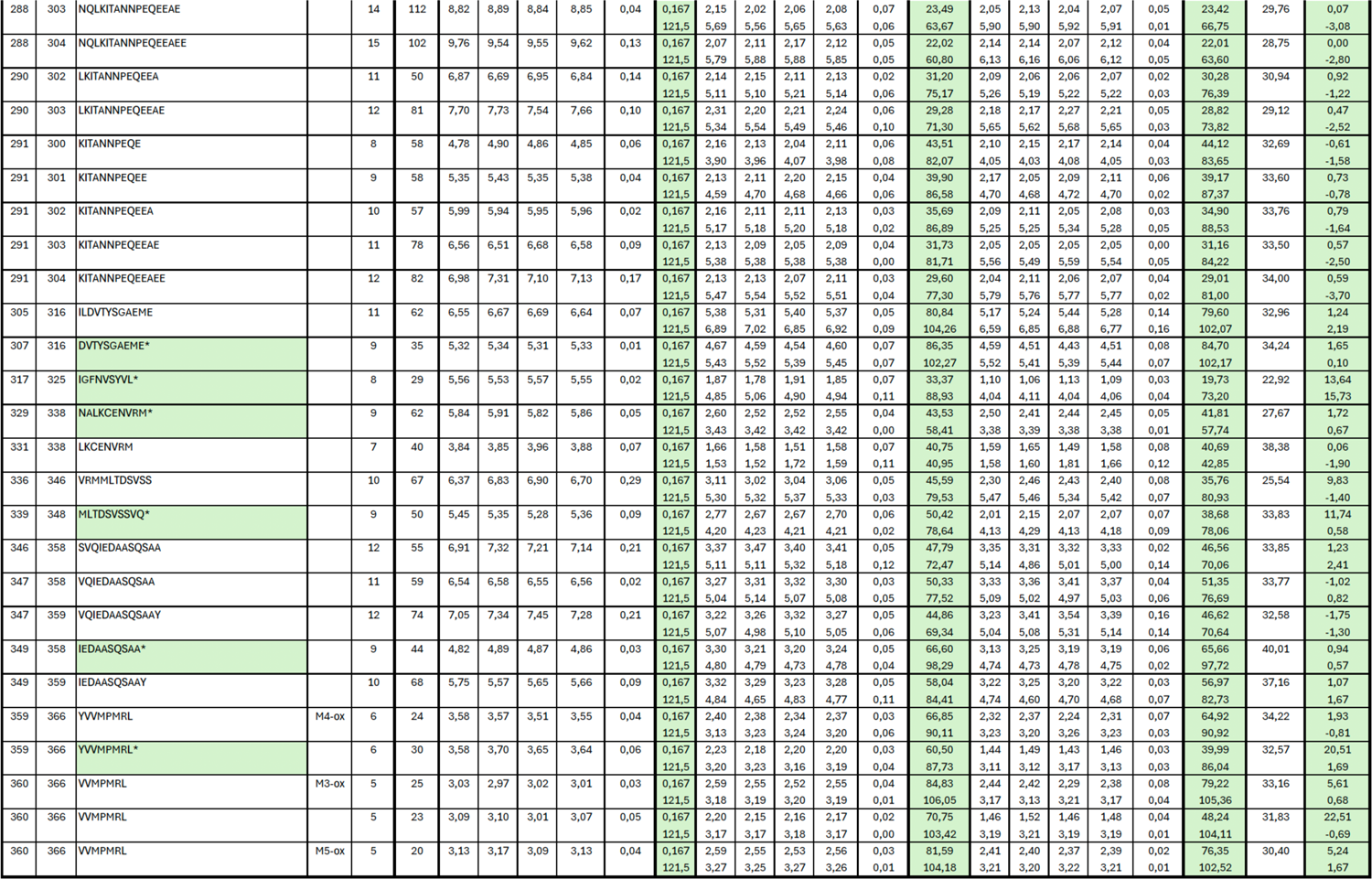
Relative deuterium uptake for all β-clamp peptides (HDX-MS)

